# Resolvin E1 improves efferocytosis and rescues severe aplastic anemia in mice

**DOI:** 10.1101/2023.02.15.528688

**Authors:** Rachel Grazda, Allison N. Seyfried, Krishna Rao Maddipatti, Gabrielle Fredman, Katherine C. MacNamara

## Abstract

Current treatments for severe aplastic anemia (SAA) rely on hematopoietic stem cell (HSC) transplantation and immunosuppressive therapies, however these treatments are not always effective. While immune-mediated destruction and inflammation are known drivers of SAA, the underlying mechanisms that lead to persistent inflammation are unknown. Using an established mouse model of SAA, we observed a significant increase in apoptotic cells within the bone marrow (BM) and demonstrate impaired efferocytosis in SAA mice, as compared to radiation controls. Single-cell transcriptomic analysis revealed heterogeneity among BM monocytes and unique populations emerged during SAA characterized by increased inflammatory signatures and significantly increased expression of *Sirpa* and *Cd47*. CD47, a “don’t eat me” signal, was increased on both live and apoptotic BM cells, concurrent with markedly increased expression of signal regulatory protein alpha (SIRPα) on monocytes. Functionally, SIRPα blockade improved cell clearance and reduced accumulation of CD47-positive apoptotic cells. Lipidomic analysis revealed a reduction in the precursors of specialized pro-resolving lipid mediators (SPMs) and increased prostaglandins in the BM during SAA, indicative of impaired inflammation resolution. Specifically, 18-HEPE, a precursor of E-series resolvins, was significantly reduced in SAA-induced mice relative to radiation controls. Treatment of SAA mice with Resolvin E1 (RvE1) improved efferocytic function, BM cellularity, platelet output, and survival. Our data suggest that impaired efferocytosis and inflammation resolution contributes to SAA progression and demonstrate that SPMs, such as RvE1, offer new and/or complementary treatments for SAA that do not rely on immune suppression.

**Key Points:**

– IFNγ impairs efferocytosis in SAA, correlating with increased SIRPa^hi^ monocytes and increased CD47 expression
– Pro-inflammatory and pro-resolving lipid mediators are imbalanced in SAA, and RvE1 treatment improved efferocytosis and disease outcomes

## Introduction

Idiopathic severe aplastic anemia (SAA) is a rare form of bone marrow failure (BMF) in which T cells drive the loss of hematopoietic stem and progenitor cells (HSPCs), resulting in pancytopenia.^1, 2^ Patients diagnosed with SAA have significantly increased interferon gamma (IFNγ) and TNF levels in the blood and BM.^3, 4^ Current therapies for SAA remain limited to immunosuppressive therapy (IST; anti-thymocyte globulin and cyclosporine) and BM transplantation.^5–7^ Although the mechanism was unknown in 1981, treatment of patients with anti-thymocyte globulin (ATG) in conjunction with BM transplantation increased survival odds immensely.^5^

Using IFNγ adenylate-uridylate–rich element (ARE)–deleted (del) mice, which exhibit increased IFNγ levels, Lin et al demonstrated that autoreactive T cells did not directly mediate destruction of HSCs during SAA, rather, IFNγ acted on the HSPC compartment to inhibit progenitor cell maturation and differentiation.^1^ It has also been shown in a murine model of SAA induced by lymphocyte transfer, that TNF produced by host macrophages drives IFNγ production and infiltration of T cells into the BM during SAA.^2^ Additionally, macrophages were shown to drive disease in an IFNγR- and CCR5- dependent manner.^8, 9^ Therefore, although T cells are an important target of current IST, additional cell types likely contribute to disease pathogenesis.

In 1995, researchers noted that patients with aplastic anemia exhibited an increased frequency of apoptotic CD34^+^ progenitor cells in the BM, and the frequency of apoptotic cells correlated with severity of disease.^10^ In addition, rampant cell death, driven by the Fas/FasL pathway for apoptosis, leads to the destruction of the hematopoietic stem and progenitor pool,^11^ as well as other cells.^12^ Macrophages are critical for SAA progression, in part through producing and augmenting inflammatory mediators,^2, 8, 9^ though macrophages are also important phagocytes within the BM where they remove apoptotic cells^13, 14^ and promote tissue regeneration.^15^

Efficient clearance of apoptotic and dead cells by phagocytes, termed efferocytosis, is crucial for inflammation-resolution and tissue function at homeostasis.^16^ Apoptotic cells provide both “find me” and “eat me” signals to phagocytes,^17^ and phosphatidylserine (PS) is a well-known “eat me” signal recognized by tissue-resident phagocytes.^18, 19^ During apoptosis the phospholipid bilayer is rearranged to expose PS on the outer leaflet of the plasma membrane, where it is either recognized directly by TAM receptors on phagocytes, or via the linker molecules Gas6 or Protein S.^17, 20^ Cell uptake and clearance is tightly regulated, however, to prevent aberrant removal of healthy cells. Canonical “don’t eat me” signals, such as CD47, interact with inhibitory receptors such as SIRPα, resulting in the inhibition of efferocytosis.^21, 22^

Resolution of inflammation is an active process mediated by a class of lipid mediators termed specialized pro-resolving mediators (SPMs).^23^ SPMs, such as the D- and E-series resolvins, derived from omega-3 polyunsaturated fatty acids, possess potent pro-resolving actions and are essential for resolution.^23^ Efferocytosis is tightly linked with the resolution of inflammation because SPMs enhance efferocytosis in a feed forward manner, further promoting the synthesis of SPMs.^16^ Resolvin E1 (RvE1) has been shown to improve efferocytosis in various inflammatory disease models of both infectious and non-infectious origins, including periodontitis,^24^ colitis,^25^ acute lung injury and bacterial pneumonia,^26^ lung cancer,^27^ asthma,^28^ and atherosclerosis.^29^ Using an established, preclinical murine model of SAA, we demonstrate dysfunctional efferocytosis and an imbalance in pro-inflammatory and pro-resolving lipid mediators. We identify a unique population of SIRPα^hi^ monocytes, and an accumulation of CD47^+^ apoptotic cells within the BM during SAA, that correlate with impaired cell clearance. RvE1 treatment not only improved efferocytosis, but also provided significant protection against disease parameters and death in mice with established SAA disease. Our findings provide new insight to the dysregulation of inflammation resolution programs in SAA pathogenesis. Moreover, our work offers key rationale to pursue novel therapeutic strategies to treat BMF that improve inflammation resolution without limiting host defense.

## Methods

### Mice

All animal protocols were approved by the Institutional Animals Care and Use Committee at Albany Medical College. C57BL/6 (H^b/b^) and BALB/cAnN (H-2^d/d^) mice were purchased from Taconic (Albany, NY, https://www.taconic.com/). Macrophage insensitive to IFNγ (MIIG) mice were generously gifted by Dr. Michael Jordan. Hybrid B6 F1 (H-2^b/d^) mice were generated by crossing C57BL/6 or MIIG mice on a C57BL/6 background with BALB/c mice. All mice were bred and housed in the Animal Resource Facility under microisolator conditions at Albany Medical College.

### Bone Marrow Failure Induction

A splenocyte-infusion method was used to induce bone marrow failure (BMF) in Hybrid F1 mice aged 6-9 weeks. Hybrid F1 mice were subjected to sub-lethal radiation (300 RADs) using a ^137^Cs source. Irradiated mice received adoptive transfer of 6.5 x 10^6^ C57BL/6 splenocytes from age- and gender-matched donors via intraperitoneal injection. Mice were euthanized via CO_2_ inhalation followed by cervical dislocation.

### Blood Collection and CBCs

Blood was collected from euthanized mice into EDTA-coated tubes via cardiac puncture and analyzed with a Heska Element HT5 for complete blood count (CBC).

### Cell Preparation and Flow Cytometry

Whole bone marrow was flushed from femurs and tibias. After RBC lysis, cell suspensions were plated and stained. See supplemental Table 1 for antibody details. Data were collected using a FACSymphony A3 (BD Biosciences) with FACSDiva software or Cytek Northern Light (Cytek Bioscienes) with SpectroFlo software and analyzed using FlowJo software (TreeStar, Ashland, OR).

### Phagocytosis Assay

200μL fluorescent Dil (DilC_18_(3))-labeled liposomes (Liposoma) were administered to mice via retro-orbital I.V. injection on day 9 post induction. BM and blood was harvested 16 hours post injection.

### Efferocytosis Assay

Whole BM was flushed and incubated with phosphatidylserine (PS)-coated lipid microparticles (Echelon Biosciences) for 1 hour at 37°C. Cells were then stained for flow cytometry.

### SIRPα Neutralization

Anti-SIRPα (clone P84; 200 μg; BioXCell) antibody was diluted in PBS and administered to mice via intraperitoneal injection on days 7 and 9 post-induction for day 10 harvest, or days 7, 9, 11, and 13 for day 14 harvest.

### Sorting and Single-Cell Analysis

Whole BM was flushed and 7-AAD^-^ CD11b^+^Ly6C^+^Ly6G^-^ cells were sorted on a BD FACSAria^TM^ Cell Sorter. Cells from each experimental group (radiation and SAA, n = 3 mice per group) were pooled together. Sample preprocessing for sequencing was performed using Chromium Next GEM Single Cell 5’ kit (10x Genomics). Sequencing and genome alignment were performed by the Center for Functional Genomics at SUNY Albany. Count matrices were loaded into R (version 4.3.1) using standard Seurat workflow. Cells with >25% mitochondrial RNA or <1000 detected genes were removed. Because the samples clustered downstream due to differences in conditions (Radiation and SAA), integration was performed utilizing the reference mapping approach described in Stuart et al.^30^ To focus on the monocytes, other cell clusters were removed from analysis.

### RvE1 Treatment

Resolvin E1 (250ng; Cayman Chemical) was diluted in PBS and administered to mice via intraperitoneal injection on days 7, 9, 11 for day 12 harvest, or days 7, 9, 11, and 13 for day 14 harvest.

### Gene Expression

Whole BM was flushed and pooled from hind limbs of three mice per group. RBCs were lysed and cells were stained to sort purified monocytes (CD11b^+^Ly6C^hi^Ly6G^-^). Monocytes were collected in RLT lysis buffer (Qiagen). mRNA was isolated (Qiagen RNeasy Mini Kit) and quantitative-RT-PCR was performed (Eppendorf *realplex*^2^ Mastercycler).

### Lipidomic analysis

Whole BM was flushed from mouse hind limbs (femurs and tibias) with ice cold PBS. Samples were flash frozen on dry ice before shipment to Wayne State University Lipidomics Core facility for analysis. Fatty acyl lipidomic analysis was performed as per published procedures.^31, 32^ Briefly, the frozen samples were thawed on ice at the time of analysis, and homogenized using Zirconium beads (Precellys, Biotage). The homogenate was extracted for fatty acyl lipids using StrataX columns (Phenomenex) following supplementation with internal standards. The extracts were analyzed by LC-MS/MS using Multiple Reaction Monitoring (MRM) method on a QTRAP5500 mass analyzer (Sciex). Identities of the individual lipid mediators were confirmed from the retention times and spectra recorded for each detected peak and were quantified relative to the internal standards. The data were normalized against protein content of the samples (ng/mg protein).

### Survival Studies

PBS or 250 ng RvE1 was administered via intraperitoneal injection days 7, 9, 11, and 13 post induction. Mice were monitored twice daily for 28 or 37 days and euthanized when moribund. Euthanasia criteria was based on signs of dehydration, response to physical stimuli, and mobility — as previously described.^9^

### Statistical Analysis

Data were analyzed with GraphPad Prism software (version 8.0, La Jolla, CA). Statistical analysis was performed using two-tailed Student’s t-test or Two-Way ANOVA with Tukey’s post-hoc analysis, and details are provided in each figure legend. Lipidomic data was analyzed using GraphPad Prism software (version 9.0, La Jolla, CA). Principal component analysis was performed using 22 lipid mediators detected across all samples from mice 3 days post-radiation for healthy (no radiation), radiation controls, and SAA-induced mice. Data were standardized and principal components were selected using Kaiser-Guttman’s rule. Heat map analysis was performed on LC-MS/MS data from radiation controls and SAA-induced mice, normalized to healthy controls.

Data Sharing Statement: All methods for flow cytometric analysis and lipidomic analysis are described in detail in the supplemental materials. Single-cell RNA sequencing data is available at GEO under accession number GSE237388. For original data please contact the authors.

## Results

### Monocytosis, cytopenias, and increased marrow cell death during SAA

Thrombocytopenia and decreased BM cellularity are well-known characteristics of severe aplastic anemia (SAA).^1, 8, 9^ Using a murine model of SAA induced by adoptive transfer of splenocytes to sub-lethally irradiated F1 mice,^8, 33, 34^ we observed a significant decrease in red blood cells (Figure 1A) and platelets (Figure 1B) by 10 days post splenocyte transfer (dpst). Mean platelet volume, which can be increased by inflammation, was significantly higher in SAA mice as compared to radiation controls (Figure 1C). Mice also exhibited striking lymphopenia, however the proportion and absolute number of circulating monocytes was significantly increased in SAA relative to controls (Figure 1D-E). Similarly, BM monocytes (CD11b^+^Ly6C^hi^Ly6G^-^) were increased by both frequency and number in SAA mice, despite extensive BM hypocellularity by 10 dpst. In contrast, neutrophils (CD11b^+^Ly6C^lo^Ly6G^hi^) were decreased (Figure 1F-H; supplemental Figure 1). Analysis of monocyte gene expression indicated an increase in *Ifnγ* and *Tnf*, known drivers of disease,^1, 2^ in SAA mice, relative to radiation controls (supplemental Figure 2). Therefore, disease progression correlated with a hematopoietic program favoring monopoiesis.

**Figure 1.**
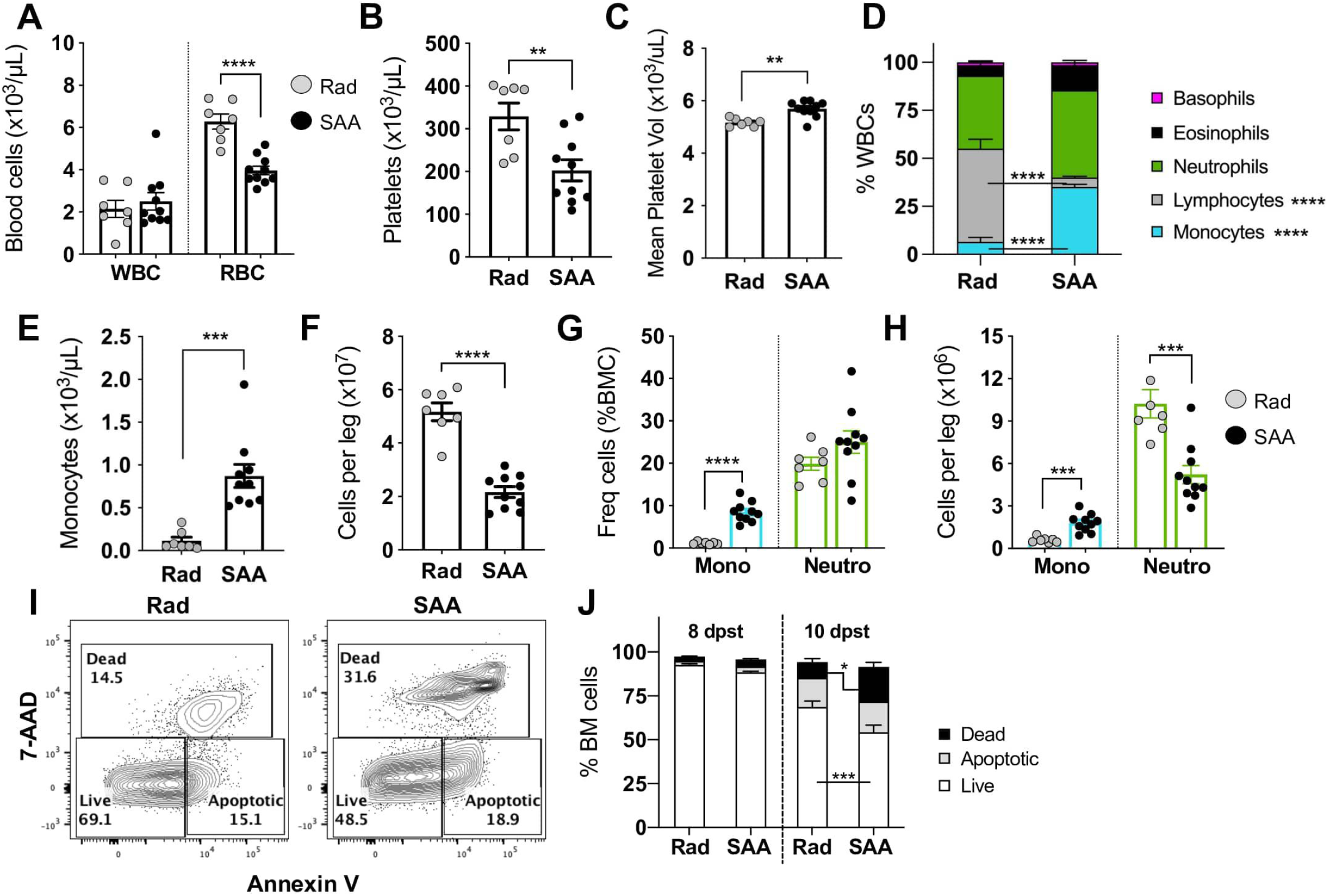
Cytopenias and BM hypocellularity are associated with increased cell death in SAA. F1 hybrid mice were induced to develop severe aplastic anemia via the radiation followed by splenocyte transfer model. Mice were euthanized 10 days post splenocyte transfer (dpst) and complete blood count data are shown for total WBCs and RBCs (**A**), platelets (**B**), and mean platelet volume (**C**). The breakdown of all WBCs (**D**) and total blood monocytes (**E**) is shown. (**F**) Overall BM cellularity in radiation control and SAA mice. (**G**) Frequency of BM monocytes (CD11b^+^Ly6C^hi^Ly6G^-^) and neutrophils (CD11b^+^ Ly6C^low^ Ly6G^+^). (**H**) Absolute number of BM monocytes and neutrophils. Data shows two pooled independent experiments showing mean ± SD, n=7-10 per group, significance was determined using a Student’s t-test. ** p<0.01, *** p<0.001, **** p<0.0001. (**G**) Gating strategy for Annexin V and 7-AAD staining to differentiate live, apoptotic, and dead BM cells, plots representative for radiation control and SAA mice. Numbers reflect the percent of cells within the gated region. (**H**) Frequency of BM cells that are live (open), apoptotic (grey), or dead (black) for both radiation controls and SAA mice days 8 and 10 post splenocyte transfer. Data shows two pooled independent experiments per time point showing mean ± SD, n=5-10 per group, significance was determined using a Student’s t-test. * p<0.05, *** p<0.001

SAA is characterized by a profound loss of BM cells, and to investigate the kinetics of cell death during disease progression we performed Annexin V and 7-AAD staining (Figure 1I). Similar frequencies of apoptotic (AnnV^+^7-AAD^-^) and dead (7-AAD^+^) cells were observed between radiation controls and SAA-induced mice at 8 dpst (Figure 1J). However, by 10 dpst SAA-induced mice had a striking increase in dead (7-AAD^+^) cells, compared to radiation controls. IFNγ can induce apoptosis^35^ and transgenic mice containing macrophages insensitive to IFNγ (referred to as MIIG mice) exhibit protection from SAA.^8^ SAA-induced MIIG mice exhibited a decrease in the frequency of dead cells in the BM, relative to littermate control (LC) counterparts (supplemental Figure 3), demonstrating that accumulation of dead cells during the early stages of SAA correlated with disease progression. These data also suggest that clearance of dead cells may be impaired in SAA.

### Single-cell transcriptomics reveals heterogeneity among BM monocytes

The hematopoietic bias towards monocyte production and accumulation of dead and dying cells suggested that impaired clearance, and defects in BM monocyte lineage cells, correlated with SAA progression. To investigate this further, we next analyzed BM monocytes by performing single-cell RNA sequencing on 7-AAD^-^ CD11b^+^ Ly6G^-^ Ly6C^+^ sorted BM cells. A total of 798 cells from radiation control and 825 cells from SAA mice were sequenced, and cells classified as monocytes were extracted from the data set (supplemental Figure 4A-C, Figure 2A). We observed three distinct monocyte clusters, of which, monocyte population 2 (Mono2) made up a majority (58%) in radiation control mice. Meanwhile, Mono2 decreased significantly during SAA (35%), while both monocyte 1 (Mono1) and monocyte 3 (Mono3) expanded (Figure 2B). Utilizing differential gene analysis, we observed that Mono1 was enriched for antigen processing/presentation (*H2-Ab1, H2-Eb1, H2-Aa, Ciita)*, while Mono3 was enriched for proliferation/cell cycle genes (*Pclaf, Stmn1, Top2a, Ccna2*; Figure 2C).

**Figure 2.**
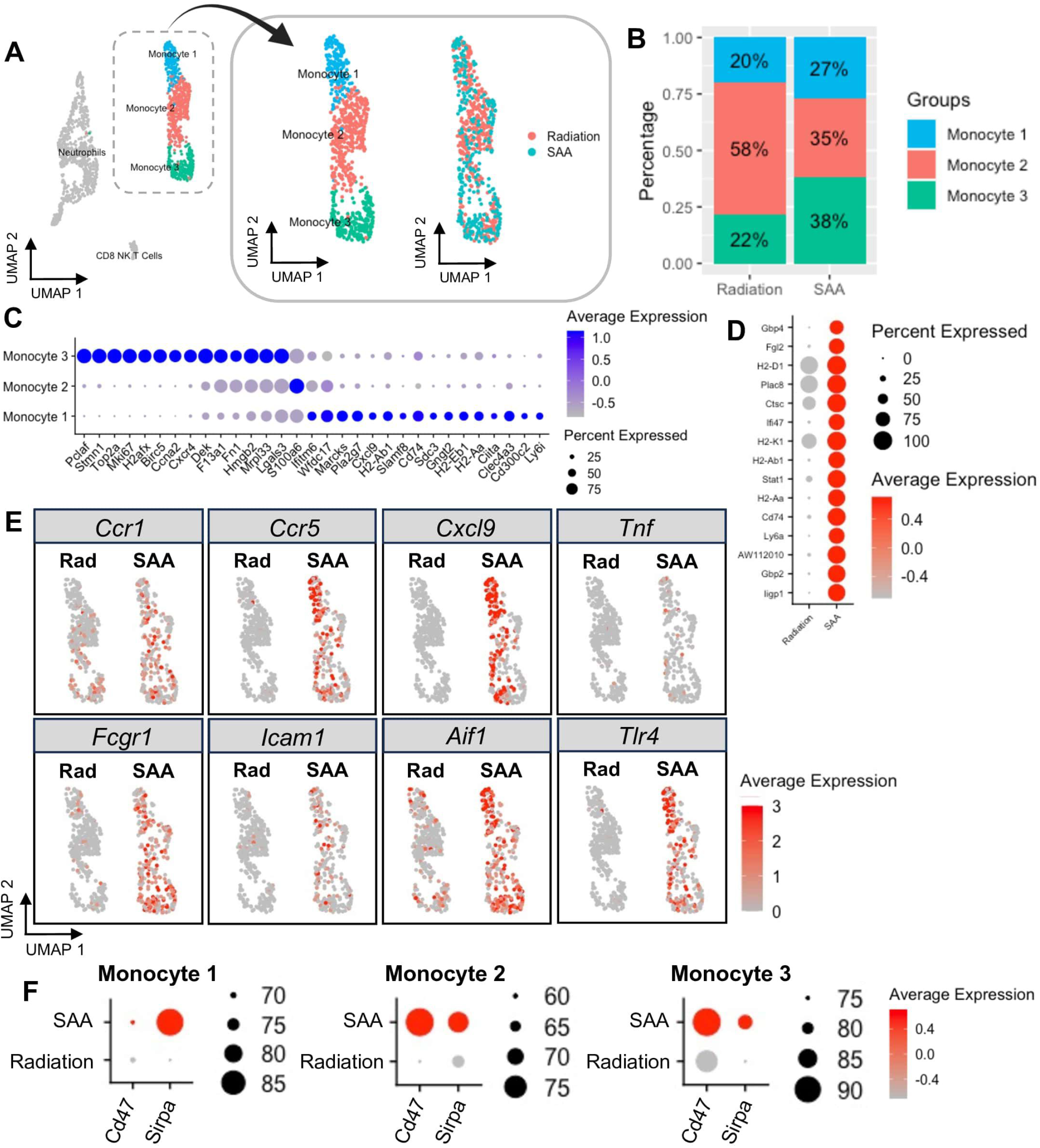
Single-cell sequencing reveals heterogeneity among BM monocytes. (**A**) UMAP plot of all cell clusters and extraction of three distinct monocyte clusters identified in BM samples from radiation control and SAA-induced mice. (**B**) Proportion and distribution of monocyte subsets in each sample. (**C**) Dot plot of the top differentially expressed genes in each monocyte population compared to the others. The size of the dot corresponds to the percentage of cells expressing each gene, while the color represents the average gene expression level. (**D**) Dot plot of top differentially expressed genes in SAA BM monocytes compared to radiation control. (**E**) Feature plot of inflammatory gene expression in radiation control and SAA. (**F**) Dot plot of *Cd47* and *Sirpa* expression on each monocyte population.

The top 15 differentially expressed genes in the SAA mice, compared to radiation control, were also examined (Figure 2D). Consistent with the role of IFNγ as a main driver of disease, monocytes from SAA mice were strongly enriched for various interferon-stimulated genes (*Fgl2, Ifi47, Stat1, Ly6a, Gbp2, Gbp4, Iigp1)*. Based on these results, and our monocyte gene expression data, other markers of inflammation were investigated. Many genes associated with inflammation, such as *Ccr1*, *Ccr5*, *Cxcl9*, *Tnf*, *Tlr4*, *Fcgr1*, *Icam1*, and *Aif1*, were highly upregulated in SAA, compared to radiation control (Figure 2E, supplementary Figure 4D). These data suggest that monocytes may perpetuate inflammation and contribute to disease progression.

Our observation that dead cells accumulated in the BM of SAA-induced mice suggested that efferocytosis may be impaired in SAA, because apoptotic cells are typically cleared rapidly.^17^ Healthy cells actively suppress their engulfment via expression of “don’t eat me” markers, such as CD47, that bind inhibitory receptors on phagocytes, including SIRPα. As cells undergo apoptosis they lose CD47 expression enabling phagocytic clearance,^36, 37^ and the CD47-SIRPα axis has been associated with impaired efferocytosis in several disease contexts.^38–40^ We therefore interrogated our single cell data set and found that all three monocyte populations significantly upregulated *Cd47* and *Sirpa* during SAA (Figure 2F). These data suggest that increased CD47 and SIRPα on monocytes may contribute to the accumulation of dead cells.

### Increased surface expression of SIRP**α** and CD47 during SAA

We next performed flow cytometry on BM myeloid cells and consistent with our scRNAseq analysis, monocytes and F4/80^+^ macrophages from SAA-induced mice exhibited a striking increase in SIRPα expression relative to radiation control mice (Figure 3A-D and supplemental Figure 5). At the same time, we observed significantly increased CD47 expression on both live and apoptotic BM cells in SAA-induced mice, relative to radiation controls (Figure 3E-G). The percentage of apoptotic BM cells expressing CD47 increased dramatically to nearly 80% in SAA-induced mice. Furthermore, we found that IFNγ was critical for SAA-induced increase in SIRPα expression among monocytes, as MIIG mice exhibited significantly reduced SIRPα^+^ monocytes and reduced MFI of SIRPα on these cells (Figure 3H-K). As MIIG mice are protected from disease, these findings support the role of aberrant CD47-SIRPα expression in disease progression. At the same time, MIIG mice had reduced expression of CD47 among BM monocytes as compared to the littermate controls under SAA conditions (Figure 3L-N). Therefore, IFNγ signaling in macrophage lineage cells promotes emergence of SIRPα^hi^ monocytes and increased CD47 expression in the BM. These findings suggest that increased CD47 on apoptotic cells, combined with increased SIRPα^hi^ monocytes, may prevent efficient clearance in the marrow during SAA.

**Figure 3.**
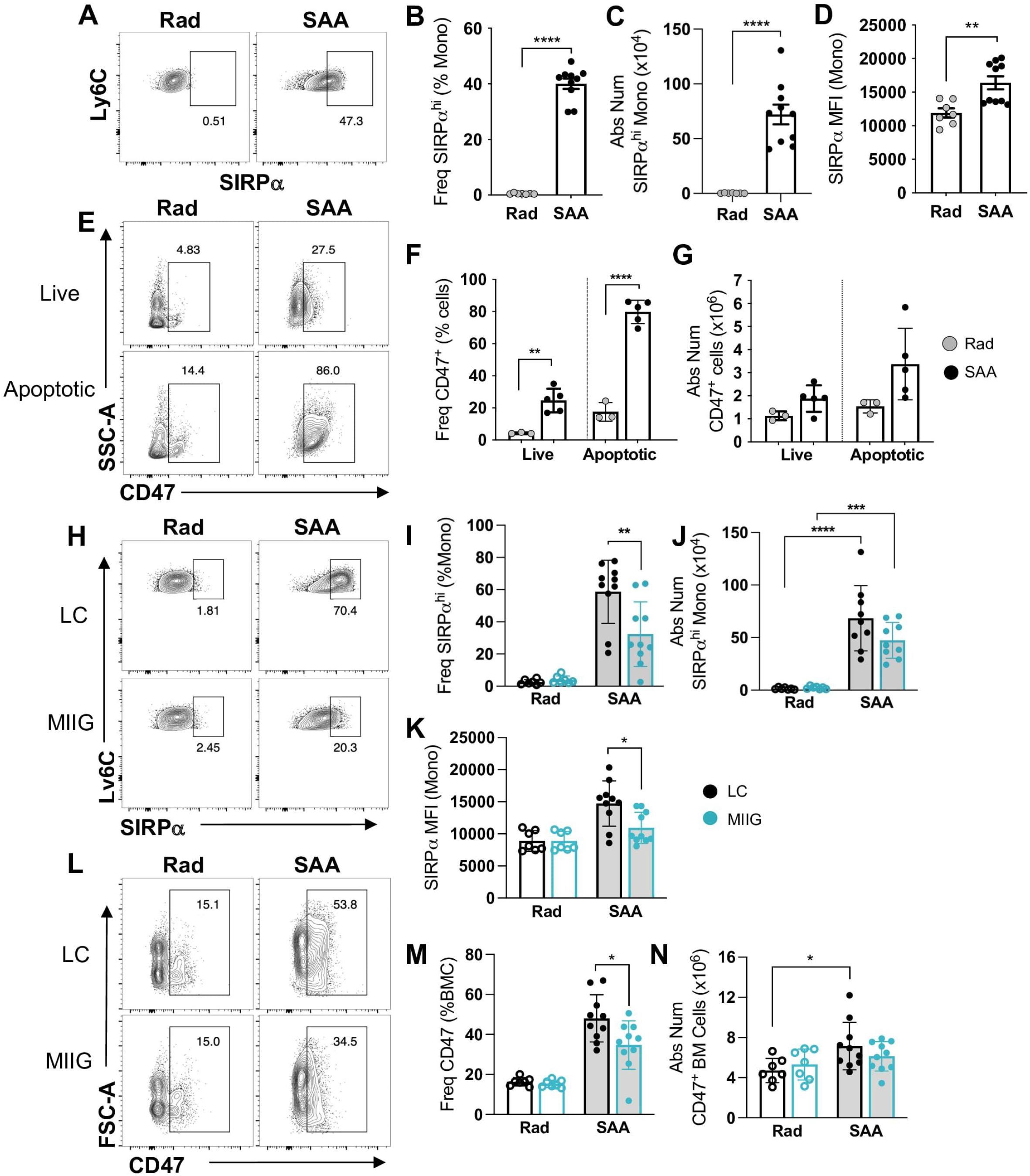
The expansion of SIRPα^hi^ monocytes and CD47 on apoptotic cells in SAA mice. Mice were induced to develop SAA and BM harvested 10 days post splenocyte transfer (dpst). (**A**) Representative gating for SIRPα^hi^ monocytes. Numbers on plots reflect the percent of cells within the gated region. The frequency, absolute number, and MFI of SIRPα^hi^ monocytes are shown in **B, C**, and **D** respectively. Data represents two pooled independent experiments showing mean ± SD, n=7-10 per group, significance using a Student’s t-test. ** p <0.01, **** p<0.0001. (**E**) Gating strategy for CD47 on live and apoptotic BM cells. Frequency (**F**) and absolute number (**G**) of CD47 for live and apoptotic cells. Data representative of one experiment showing mean ± SD, n=3-5 per group. Significance was determined using a Two-way ANOVA with Tukey’s multiple comparison test. * p<0.05, ** p <0.01, *** p<0.001, **** p<0.0001.

### SIRP**α**-CD47 axis contributes to disease progression

To test whether the SIRPα−CD47 axis could be targeted therapeutically, SAA-induced mice were treated with anti-SIRPα neutralizing antibody days 7, 9, and 11 post induction (supplementary Figure 6A). We observed an increase in BM cellularity and a significant decrease in the frequency of dead cells and CD47^+^ apoptotic cells with anti- SIRPα treatment (supplementary Figure 6B-E), thus supporting the notion that SIRPα limits efficient efferocytosis. However, as expected, anti-SIRPα treatment also exacerbated anemia during SAA by 14 dpst (supplementary Figure 6F-I), consistent with the role of CD47 as an important “marker of self” on healthy cells, especially red blood cells.^41^ Therefore, while these data support the idea that the SIRPα−CD47 axis contributes to the accumulation of dead cells during SAA, this axis is necessary for RBC circulation precluding its use as a treatment for SAA.

### Decreased phagocytosis and efferocytosis in SAA

To directly examine whether phagocytes were functionally impaired during SAA, F1 mice were induced to develop disease and administered fluorescently-labeled (Dil-labeled) liposomes day 9 post-induction to evaluate phagocytotic capacity. BM was harvested day 10 to evaluate the uptake of liposomes via flow cytometry. SAA mice exhibited a decrease in Dil liposome-positive and Dil MFI on monocytes and F4/80^+^ macrophages, suggesting impairments in phagocytosis (supplemental Figure 7).

To more specifically address whether efferocytosis was defective during SAA, we utilized phosphatidylserine (PS)-coated microparticles to mimic apoptotic cells in an *ex vivo* assay. BM from SAA-induced and radiation control mice was incubated with fluorescently tagged PS-coated lipid microparticles (PS-MPs), and uptake was determined via flow cytometry (Figure 4A-B). We observed a significant reduction in PS- MP^+^ monocytes and F4/80^+^ macrophages in marrow isolated from SAA-induced mice, relative to controls, at 8 dpst (Figure 4C-E). Consistent with the role of impaired efferocytosis in disease progression, SAA-induced MIIG mice exhibited no change in efferocytosis whereas LC mice had significantly reduced uptake of PS-MPs in SAA conditions (Figure 4F-I). These data demonstrate that during SAA progression unique inflammatory monocytes emerge in the BM and exhibit functional impairments.

**Figure 4.**
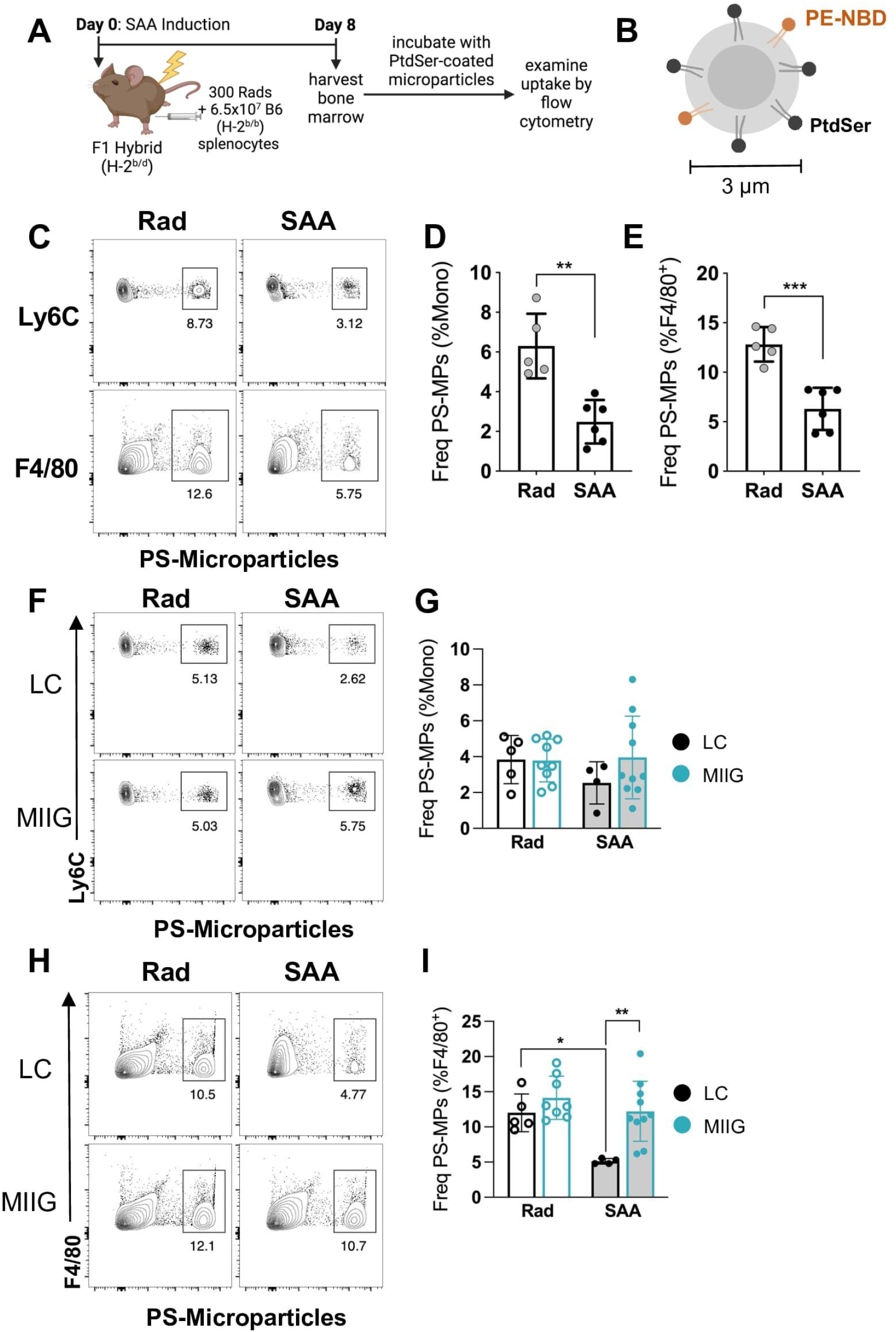
Enhanced SIRPα and CD47 expression is associated with impaired efferocytosis. (**A**) F1 hybrid mice were induced with SAA and BM harvested 8 dpst. Whole BM was flushed and incubated with PS lipid microparticles (PS-MPs) for 1 hour at 37°C. (**B**) The lipid microparticles contain attached PS with the phospholipid headgroup exposed and trace amounts of a fluorescent lipid PE-NBD embedded in the particles. (**C**) Representative staining of monocytes and F4/80^+^cells. Numbers on plots reflect the percent of cells within the gated region. Frequencies of PS-MPs-positive cells among total monocytes (**D**) and F4/80^+^ macrophages (**E**). Data representative of one experiment showing mean ± SD, n=5-6 mice per group. Significance was determined using a Student’s t-test. Representative staining (**F**) and frequency (**G**) of PS-MP^+^ monocytes from LC and MIIG mice. (**H**) Representative staining of PS-MP^+^ F4/80^+^ macrophages from LC and MIIG mice. (**I**) Frequencies of PS-MP^+^ cells among total F4/80^+^ BM cells. Data representative of two independent experiments showing mean ± SD, n=4-10 mice per group. Significance was determined using a Two-way ANOVA with Tukey’s multiple comparison test. * p<0.05, ** p<0.01, *** p<0.001.

### Imbalance of lipid mediators that regulate inflammation resolution in SAA

We next questioned whether key lipid mediators associated with inflammation resolution programs were impacted during disease progression. To do this we took an unbiased approach and evaluated the polyunsaturated fatty acid (PUFA) metabolome in BM of healthy, radiation control, and SAA-induced mice via LC-MS/MS. We performed principal component analysis (PCA) which revealed that a majority of the variation (PC1; 43%) at day 3 post-radiation was due to differences in mono-hydroxy eicosapentaenoic acids, including a decrease in 18-HEPE in SAA mice, relative to radiation controls (Figure 5A, supplemental Figure 8A). Indeed, limited 18-HEPE, the precursor to the E-series resolvins, may contribute to impaired generation of these SPMs in the marrow during SAA (Figure 5B). By day 8, SAA-induced mice had significantly elevated prostaglandins (PGE2 and PGD2) and TXB2 relative to radiation controls (Figure 5C-E, supplemental Figure 8B, supplementary Table 2). Our data demonstrate that in the context of SAA, omega-3 derived lipid precursors are decreased in the marrow during initiation of disease, whereas progression correlates with unregulated prostanoid and thromboxane synthesis. Together, these data support the conclusion that inflammation resolution kinetics are defective in SAA.

**Figure 5.**
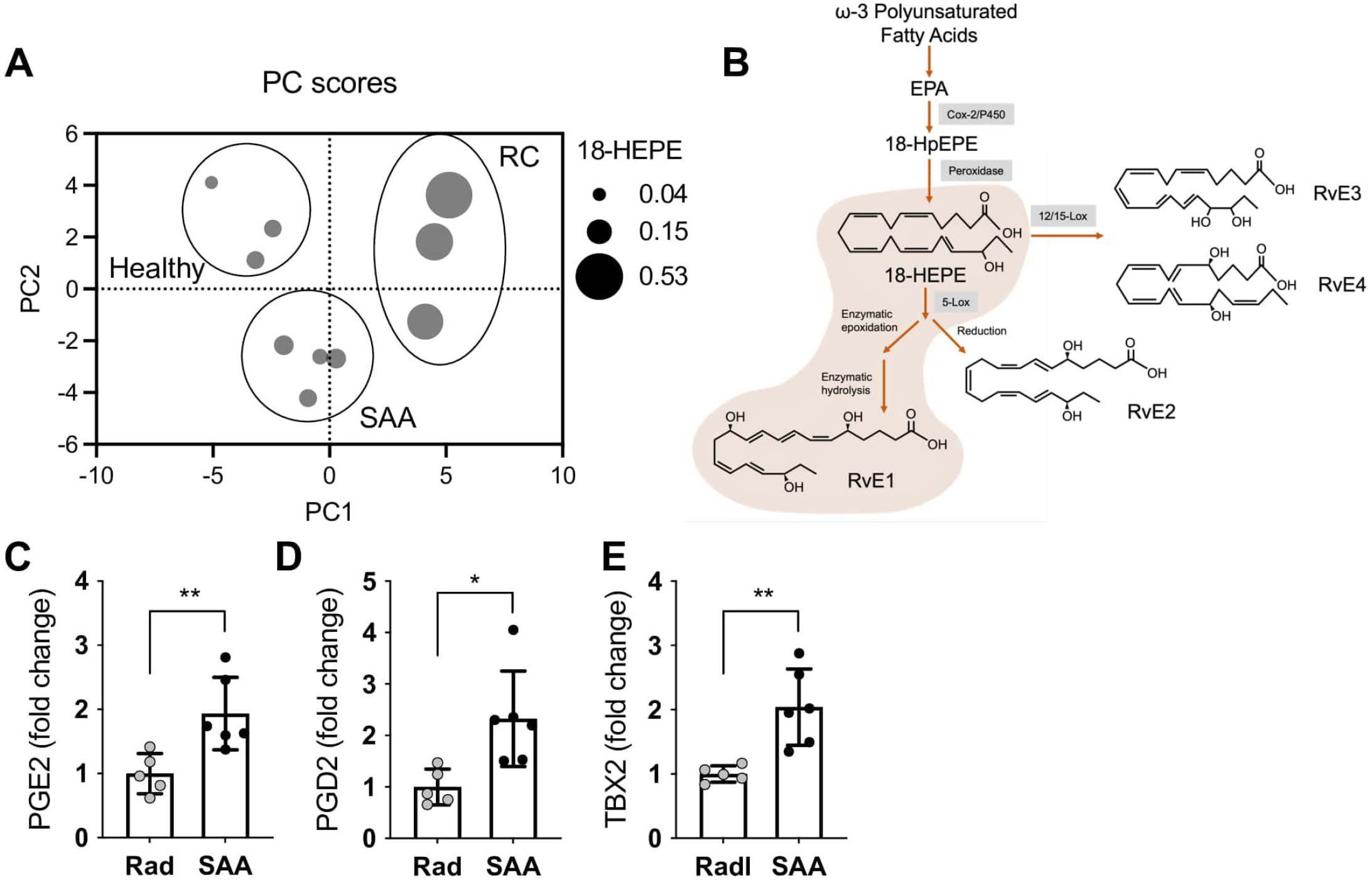
Imbalanced pro-Inflammatory- pro-resolving lipid mediators. Bone marrow was collected 3 or 8 dpst and analyzed by LC-MS/MS. (**A**) Principal component analysis of samples taken from mice without exposure to radiation (healthy), radiation only (RC), or radiation plus splenocytes to induce SAA (SAA) day 3 post-radiation. Each dot represents data from a single mouse and the size of the dot reflects concentration of 18-HEPE. (**B**) Schematic of the biosynthesis of resolvins from ω-3 polyunsaturated fatty acids. Fold change in concentrations of PGE2 (**C**), PGD2 (**D**), or TXB2 (**E**) in the bone marrow of SAA mice, relative to radiation control, 8 dpst. Data representative of two independent experiments that included 2-3 mice each. Significance was determined using a Student’s t-test. * p<0.05, ** p<0.01.

### Resolvin E1 provides therapeutic benefit in SAA

To address whether SPMs may be efficacious in treating SAA, we first examined expression of key SPM receptors on BM monocytes. We determined by flow cytometry that ChemR23 (*Cmklr1*), the receptor for E-series resolvins,^42^ was significantly upregulated on BM monocytes, neutrophils, and T cells, but not on macrophages or LK cells (supplemental Figure 9A-E). Gene expression analysis from our single-cell dataset demonstrated an increase in *Cmklr1* during SAA, but especially on the Mono1 subset, which exhibited the greatest *Sirpa* expression (supplemental Figure 9F). Together, these data provided rationale for testing our hypothesis that exogenous RvE1 would improve efferocytosis and mitigate SAA progression.

First, to examine whether exogenous RvE1 could improve efferocytosis in the context of SAA, we administered RvE1 or vehicle every other day, beginning at day 7 post-induction, a timepoint when the marrow is already hypocellular (Figure 6A). We observed that RvE1 treatment decreased expression of the inhibitory receptor SIRPα on both monocytes and macrophages in the BM (Figure 6B-C). The RvE1-induced reduction in SIRPa expression correlated with increased uptake of PS-MPs with a significant increase in the frequency of PS-MP^+^ monocytes at day 12 dpst (Figure 6D- E). Although we observed a moderate increase in PS-MP^+^ F4/80^+^ cells by 12 dpst this did not reach statistical significance (Figure 6F-G). These data support the conclusion that RvE1 improves monocyte efferocytic function in SAA-induced mice RvE1 treatment improved WBC count and platelets by 14 dpst (Figure 7A-C). While RvE1 did not improve lymphopenia, anemia, and only mildly limited SAA-induced monocytosis (Figure 7D-E), RvE1 treatment increased overall BM cellularity by day 14 (Figure 7F). Total LK cells and HSC progenitor populations were also significantly increased with RvE1 treatment (Figure 7G; supplementary Figure 10). Moreover, RvE1 significantly improved survival when mice were administered therapeutic doses of RvE1, on days 7, 9, 11, and 13 post-SAA induction (Figure 7H). Together, our data demonstrate that RvE1 therapy improves efferocytic function, which correlated with protection against SAA-induced mortality. These findings suggest that dysfunctional inflammation resolution is a novel therapeutic target for improving treatments for SAA patients.

**Figure 6.**
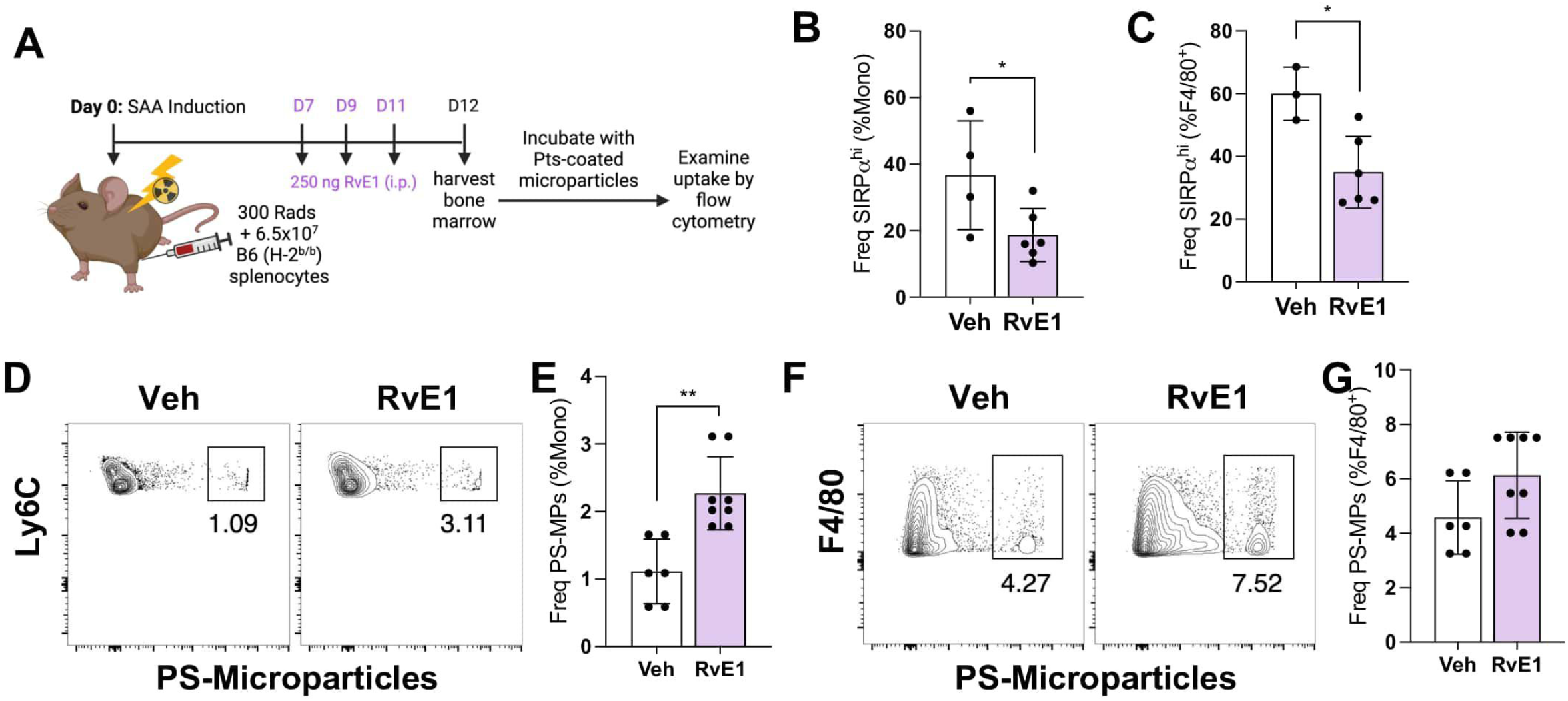
Exogenous RvE1 improves aberrant monocyte phenotype and efferocytosis. (**A**) Mice were induced to develop SAA and treated with 250ng RvE1 days 7, 9, and 11 post induction. BM was harvest day 12 dpst. The frequency of SIRPα on BM monocytes (**B**) and F4/80+ macrophages (**C**). Data representative of one experiment showing mean ± SD, n=4-6 mice per group. Staining (**D** and **F**) and frequency (**E** and **G**) of PS-MPs on monocytes and F4/80^+^ cells. Numbers on flow plots are the frequency of cells within the gated region. Data from two independent experiments with mean ± SD, n=6-8 per group. Significance was determined using a Student’s t-test. * p<0.05, ** p<0.01

**Figure 7.**
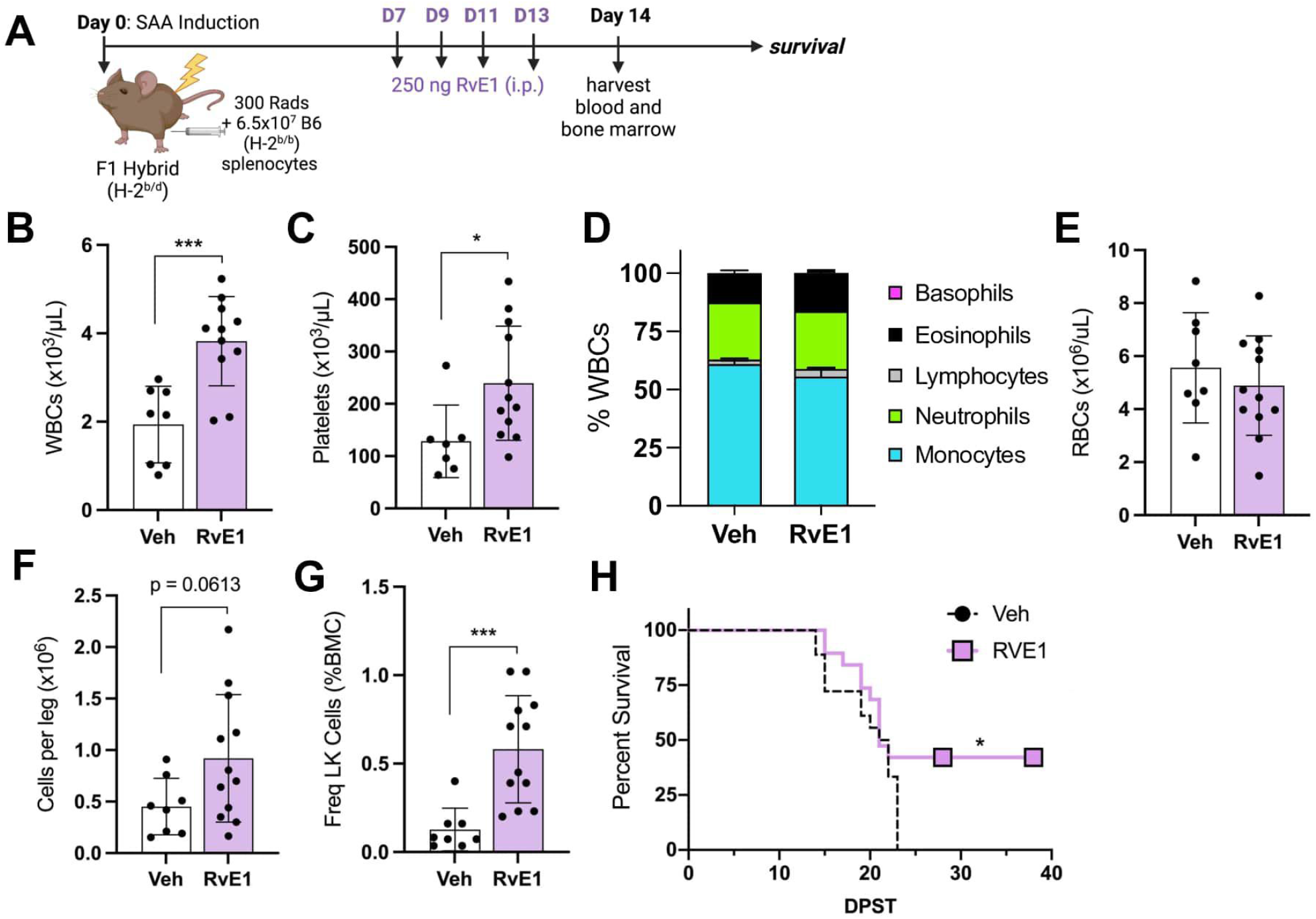
Exogenous RvE1 improves platelets, BM cellularity and survival. (**A**) SAA-induced F1 hybrid mice were treated with 250ng of RvE1 days 7, 9, 11, and 13 post induction. BM and blood was evaluated day 14. The frequency of all WBCs (**B**), platelets (**C**), WBC breakdown (**D**), and total RBCs (**C**) are shown. Total BM cellularity (**F**) and frequency of LK cells (**G**) in F1 mice treated with vehicle or RvE1. Data representative of two independent experiments showing mean ± SD, n=8-12 per group. Significance was determined using a Student’s t-test. * p<0.05, *** p<0.001. (**I**) Percent survival 37 days post-induction; SAA mice treated with vehicle (black dashed line) *n = 18*; SAA mice treated with RvE1 (purple line) *n = 19*. Data pooled from three independent experiments.

## Discussion

Current treatments for SAA rely on HSC transplantation, when possible, and IST, though IST has a high refractory rate. Recently approved Etrombopag, an agonist of the thrombopoietin receptor, has shown improved response times in combination with IST, although overall responses were not improved in older patients or those that began treatment with severe disease.^58^ Recent preclinical studies using the Janus kinase (Jak1/2) inhibitor Ruxolitinib (Rux) demonstrated effectiveness in a murine model of SAA with markedly reduced T lymphocytes in the marrow.^59^ However, toxicity associated with Rux use has been noted in humans and mouse models.^60, 61^ Therefore, improved therapeutic strategies are still needed for durable treatment of SAA. While suppressing inflammation is an important target for treating disease in SAA, we reasoned that understanding mechanisms underlying non-resolving inflammation may reveal additional, novel targets for treating disease.

Efferocytosis is a crucial component of resolution pathways and maintains tissue homeostasis by limiting inflammation. By engaging in specific ligand-receptor signaling, efferocytes promote the resolution of inflammation by clearing dying cells, releasing anti-inflammatory mediators and driving SPM biosynthesis.^20, 43^ Apoptotic cells that are not efficiently cleared enter secondary necrosis, characterized by the breakdown of the cellular membrane and the leakage of inflammatory DAMPs that exacerbate inflammation.^20^ Although impairments in resolution are known to underly several chronic inflammatory diseases,^44–47^ whether resolution responses are impaired during SAA or in other BMF diseases has yet to be investigated.

SAA is associated with increased apoptosis,^10^ however the accumulation of necrotic cells also suggested deficiencies in cell clearance. The increase in dead and dying cells correlated with the expansion of a unique population of SIRPα^hi^ monocytes, not observed in radiation controls. Indeed, IFNγ is known to drive monopoiesis,^48^ and previous studies demonstrated that increased monocytes preceded the accumulation of macrophages in the bone marrow during SAA pathogenesis.^8^ Consistent with a potential role of SIRPα^hi^ monocytes in driving disease, we noted that this population was significantly reduced in the MIIG mouse model, a strain that is protected from developing SAA.^8^ We reasoned that increased expression of the inhibitory receptor SIRPα impaired the clearance of apoptotic and dead cells.

To prevent improper engulfment by phagocytes, healthy cells utilize “don’t eat me” markers, such as CD47. Dying cells typically lose or alter the expression of CD47 to enable cell clearance, as in the case of aging RBCs, where CD47 is expressed on normal RBCs, preventing their elimination by phagocytes.^41^ Therefore, it is of no surprise that anti-SIRPα therapy worsened anemia in SAA-induced mice. High CD47 expression on dying cells has also been shown to promote inefficient clearance via the “nibbling” of cells, rather than whole-cell engulfment by phagocytes.^44^ TNF and IFNγ, which are increased during SAA,^2^ have been shown to upregulate CD47 expression.^38, 49^ As defective efferocytosis is known to elicit pro-inflammatory cytokines, it is possible that CD47, TNF, and IFNγ play a critical role in the feedback loop that inhibits cellular clearance and inflammation-resolution, thus potentiating inflammation.

Specialized pro-resolving mediators (SPMs) actively drive the resolution of inflammation and a return to homeostasis.^23^ Class switching of lipid mediators, from pro-inflammatory to pro-resolving, is essential for resolution to occur.^50^ Lipid mediator analysis of the BM from SAA-induced mice demonstrated an imbalance of pro-inflammatory to pro-resolving mediators with exuberant prostanoid synthesis. SPM biosynthesis occurs via transcellular mechanisms that require close contact of cells expressing distinct enzymes, and it is notable that during SAA the bone marrow becomes increasingly hypocellular, which may prevent SPM generation. A loss of proper SPM biosynthesis or signaling has been linked with the exacerbation of inflammation in various disease settings.^51–53^ While the cause is still unclear, a lack of SPM production and an increase in prostanoid synthesis, combined with dysregulated unalamation,^54^ appears to be contributing to persistent inflammation observed during SAA. A recent report uncovered a prostanoid storm limited efferocytosis in vitro and in atherosclerotic plaques,^55^ thus supporting the notion that sustained elevated prostanoids, without a compensatory increase in SPMs, may contribute to defective efferocytosis in the bone marrow during SAA.

Our findings that efferocytosis was impaired in SAA, and that SPM therapy was efficacious, provided strong rationale for targeting inflammation-resolution during BM failure. Treatment with exogenous SPMs has been utilized to improve resolution in multiple experimental settings.^25, 46, 56^ SPMs have a unique safety profile since they, unlike immunosuppressive therapies, do not compromise the host’s natural immune response.^26, 57^ During SAA, 18-HEPE, the precursor for the E-series resolvins is significantly reduced, relative to radiation controls. Meanwhile, the receptor for RvE1, ChemR23, was increased on monocytes and neutrophils in SAA. Together, these data suggested that RvE1 could be an ideal therapeutic for treating SAA. Indeed, RvE1 therapy, starting at a timepoint where the BM is already hypocellular, improved platelet count, BM cellularity, survival, and efferocytosis. Therapies aimed at improving resolution, perhaps in conjunction with lower dosing of IST, may provide a more effective and safe treatment for SAA that does not compromise the patient’s natural immune response. Moreover, improving resolution may contribute to more durable responses to IST that improve long-term outcomes.

## Acknowledgements

The authors would like to thank Jesse Bonin and Ramon Bossardi Ramos for technical assistance. This work was supported by CDMRP Bone Marrow Failure Research Program-IDA (BM190079) to K.C.M. This study was also supported in part by National Center for Research Resources, National Institutes of Health Grant S10RR027926 (K.R.M.)

## Authorship

Contribution: R.G. and A.N.S. performed *in vivo* experiments, analyzed data, and prepared figures. R.G. wrote the manuscript. K.R.M. provided oversight and guidance for LC-MS/MS studies and analyzed data. G.F. contributed to experimental design and data analysis for the project. K.C.M. conceived of the project, analyzed data, and wrote the manuscript. The order of the first authors was determined on the basis of the effort and contributions to writing of the manuscript.

Conflict-of-interest disclosure: The authors declare no competing financial interests. Correspondence: Katherine C. MacNamara, Department of Immunology and Microbial Disease, Albany Medical College, MC-151, 47 New Scotland Avenue, Albany, NY 12208.

## References

1. Lin FC, Karwan M, Saleh B, et al. IFN-gamma causes aplastic anemia by altering hematopoietic stem/progenitor cell composition and disrupting lineage differentiation. Blood. 2014;124(25):3699–3708.

2. Sun W, Wu Z, Lin Z, et al. Macrophage TNF-alpha licenses donor T cells in murine bone marrow failure and can be implicated in human aplastic anemia. Blood. 2018;132(26):2730–2743.

3. Zoumbos NC, Gascon P, Djeu JY, Young NS. Interferon is a mediator of hematopoietic suppression in aplastic anemia in vitro and possibly in vivo. Proc Natl Acad Sci U S A. 1985;82(1):188–192.

4. Merli P, Quintarelli C, Strocchio L, Locatelli F. The role of interferon-gamma and its signaling pathway in pediatric hematological disorders. Pediatr Blood Cancer. 2021;68(4):e28900.

5. Gratwohl A, Osterwalder B, Nissen C, et al. [Treatment of severe aplastic anemia]. Schweiz Med Wochenschr. 1981;111(41):1520–1522.

6. Xu L, Fu B, Wang W, et al. Haploidentical hematopoietic cell transplantation for severe acquired aplastic anemia: a case-control study of post-transplant cyclophosphamide included regimen vs. anti-thymocyte globulin & colony-stimulating factor-based regimen. Sci China Life Sci. 2020;63(6):940–942.

7. Rosenfeld S, Follmann D, Nunez O, Young NS. Antithymocyte globulin and cyclosporine for severe aplastic anemia: association between hematologic response and long-term outcome. JAMA. 2003;289(9):1130–1135.

8. McCabe A, Smith JNP, Costello A, Maloney J, Katikaneni D, MacNamara KC. Hematopoietic stem cell loss and hematopoietic failure in severe aplastic anemia is driven by macrophages and aberrant podoplanin expression. Haematologica. 2018;103(9):1451–1461.

9. Seyfried AN, McCabe A, Smith JNP, Calvi LM, MacNamara KC. CCR5 maintains macrophages in the bone marrow and drives hematopoietic failure in a mouse model of severe aplastic anemia. Leukemia. 2021;35(11):3139–3151.

10. Philpott NJ, Scopes J, Marsh JC, Gordon-Smith EC, Gibson FM. Increased apoptosis in aplastic anemia bone marrow progenitor cells: possible pathophysiologic significance. Exp Hematol. 1995;23(14):1642–1648.

11. Liu CY, Fu R, Wang HQ, et al. Fas/FasL in the immune pathogenesis of severe aplastic anemia. Genet Mol Res. 2014;13(2):4083–4088.

12. Gupta P, Niehans GA, LeRoy SC, et al. Fas ligand expression in the bone marrow in myelodysplastic syndromes correlates with FAB subtype and anemia, and predicts survival. Leukemia. 1999;13(1):44–53.

13. Michalski MN, Koh AJ, Weidner S, Roca H, McCauley LK. Modulation of Osteoblastic Cell Efferocytosis by Bone Marrow Macrophages. J Cell Biochem. 2016;117(12):2697–2706.

14. McCauley LK, Dalli J, Koh AJ, Chiang N, Serhan CN. Cutting edge: Parathyroid hormone facilitates macrophage efferocytosis in bone marrow via proresolving mediators resolvin D1 and resolvin D2. J Immunol. 2014;193(1):26–29.

15. Wynn TA, Vannella KM. Macrophages in Tissue Repair, Regeneration, and Fibrosis. Immunity. 2016;44(3):450–462.

16. Doran AC, Yurdagul A, Jr., Tabas I. Efferocytosis in health and disease. Nat Rev Immunol. 2020;20(4):254–267.

17. Elliott MR, Koster KM, Murphy PS. Efferocytosis Signaling in the Regulation of Macrophage Inflammatory Responses. J Immunol. 2017;198(4):1387–1394.

18. Shiratsuchi A, Osada S, Kanazawa S, Nakanishi Y. Essential role of phosphatidylserine externalization in apoptosing cell phagocytosis by macrophages. Biochem Biophys Res Commun. 1998;246(2):549–555.

19. Fadok VA, Voelker DR, Campbell PA, Cohen JJ, Bratton DL, Henson PM. Exposure of phosphatidylserine on the surface of apoptotic lymphocytes triggers specific recognition and removal by macrophages. J Immunol. 1992;148(7):2207–2216.

20. Korns D, Frasch SC, Fernandez-Boyanapalli R, Henson PM, Bratton DL. Modulation of macrophage efferocytosis in inflammation. Front Immunol. 2011;2(57.

21. Gardai SJ, McPhillips KA, Frasch SC, et al. Cell-surface calreticulin initiates clearance of viable or apoptotic cells through trans-activation of LRP on the phagocyte. Cell. 2005;123(2):321–334.

22. Okazawa H, Motegi S, Ohyama N, et al. Negative regulation of phagocytosis in macrophages by the CD47-SHPS-1 system. J Immunol. 2005;174(4):2004–2011.

23. Serhan CN, Levy BD. Resolvins in inflammation: emergence of the pro-resolving superfamily of mediators. J Clin Invest. 2018;128(7):2657–2669.

24. Hasturk H, Kantarci A, Ohira T, et al. RvE1 protects from local inflammation and osteoclast- mediated bone destruction in periodontitis. FASEB J. 2006;20(2):401–403.

25. Ishida T, Yoshida M, Arita M, et al. Resolvin E1, an endogenous lipid mediator derived from eicosapentaenoic acid, prevents dextran sulfate sodium-induced colitis. Inflamm Bowel Dis. 2010;16(1):87–95.

26. Seki H, Fukunaga K, Arita M, et al. The anti-inflammatory and proresolving mediator resolvin E1 protects mice from bacterial pneumonia and acute lung injury. J Immunol. 2010;184(2):836–843.

27. Kantarci A, Kansal S, Hasturk H, Stephens D, Van Dyke TE. Resolvin E1 Reduces Tumor Growth in a Xenograft Model of Lung Cancer. Am J Pathol. 2022;192(10):1470–1484.

28. Haworth O, Cernadas M, Yang R, Serhan CN, Levy BD. Resolvin E1 regulates interleukin 23, interferon-gamma and lipoxin A4 to promote the resolution of allergic airway inflammation. Nat Immunol. 2008;9(8):873–879.

29. Hasturk H, Abdallah R, Kantarci A, et al. Resolvin E1 (RvE1) Attenuates Atherosclerotic Plaque Formation in Diet and Inflammation-Induced Atherogenesis. Arterioscler Thromb Vasc Biol. 2015;35(5):1123–1133.

30. Stuart T, Butler A, Hoffman P, et al. Comprehensive Integration of Single-Cell Data. Cell. 2019;177(7):1888–1902 e1821.

31. Norris PC, Skulas-Ray AC, Riley I, et al. Identification of specialized pro- resolving mediator clusters from healthy adults after intravenous low-dose endotoxin and omega-3 supplementation: a methodological validation. Sci Rep. 2018;8(1):18050.

32. Maddipati KR, Romero R, Chaiworapongsa T, et al. Eicosanomic profiling reveals dominance of the epoxygenase pathway in human amniotic fluid at term in spontaneous labor. FASEB J. 2014;28(11):4835–4846.

33. Chen J, Lipovsky K, Ellison FM, Calado RT, Young NS. Bystander destruction of hematopoietic progenitor and stem cells in a mouse model of infusion-induced bone marrow failure. Blood. 2004;104(6):1671–1678.

34. Bloom ML, Wolk AG, Simon-Stoos KL, Bard JS, Chen J, Young NS. A mouse model of lymphocyte infusion-induced bone marrow failure. Exp Hematol. 2004;32(12):1163–1172.

35. Ossina NK, Cannas A, Powers VC, et al. Interferon-gamma modulates a p53- independent apoptotic pathway and apoptosis-related gene expression. J Biol Chem. 1997;272(26):16351–16357.

36. Lv Z, Bian Z, Shi L, et al. Loss of Cell Surface CD47 Clustering Formation and Binding Avidity to SIRPalpha Facilitate Apoptotic Cell Clearance by Macrophages. J Immunol. 2015;195(2):661–671.

37. Burger P, Hilarius-Stokman P, de Korte D, van den Berg TK, van Bruggen R. CD47 functions as a molecular switch for erythrocyte phagocytosis. Blood. 2012;119(23):5512–5521.

38. Kojima Y, Volkmer JP, McKenna K, et al. CD47-blocking antibodies restore phagocytosis and prevent atherosclerosis. Nature. 2016;536(7614):86-90.

39. Meier LA, Faragher JL, Osinski V, et al. CD47 Promotes Autoimmune Valvular Carditis by Impairing Macrophage Efferocytosis and Enhancing Cytokine Production. J Immunol. 2022;208(12):2643–2651.

40. Barrera L, Montes-Servin E, Hernandez-Martinez JM, et al. CD47 overexpression is associated with decreased neutrophil apoptosis/phagocytosis and poor prognosis in non-small-cell lung cancer patients. Br J Cancer. 2017;117(3):385–397.

41. Oldenborg PA, Zheleznyak A, Fang YF, Lagenaur CF, Gresham HD, Lindberg FP. Role of CD47 as a marker of self on red blood cells. Science. 2000;288(5473):2051-2054.

42. Ohira T, Arita M, Omori K, Recchiuti A, Van Dyke TE, Serhan CN. Resolvin E1 receptor activation signals phosphorylation and phagocytosis. J Biol Chem. 2010;285(5):3451–3461.

43. Cai B, Thorp EB, Doran AC, et al. MerTK cleavage limits proresolving mediator biosynthesis and exacerbates tissue inflammation. Proc Natl Acad Sci U S A. 2016;113(23):6526–6531.

44. Gerlach BD, Marinello M, Heinz J, et al. Resolvin D1 promotes the targeting and clearance of necroptotic cells. Cell Death Differ. 2020;27(2):525–539.

45. Rymut N, Heinz J, Sadhu S, et al. Resolvin D1 promotes efferocytosis in aging by limiting senescent cell-induced MerTK cleavage. FASEB J. 2020;34(1):597–609.

46. El Kebir D, Gjorstrup P, Filep JG. Resolvin E1 promotes phagocytosis-induced neutrophil apoptosis and accelerates resolution of pulmonary inflammation. Proc Natl Acad Sci U S A. 2012;109(37):14983–14988.

47. Martin-Rodriguez O, Gauthier T, Bonnefoy F, et al. Pro-Resolving Factors Released by Macrophages After Efferocytosis Promote Mucosal Wound Healing in Inflammatory Bowel Disease. Front Immunol. 2021;12(754475.

48. de Bruin AM, Libregts SF, Valkhof M, Boon L, Touw IP, Nolte MA. IFNgamma induces monopoiesis and inhibits neutrophil development during inflammation. Blood. 2012;119(6):1543–1554.

49. Ye ZH, Jiang XM, Huang MY, et al. Regulation of CD47 expression by interferon-gamma in cancer cells. Transl Oncol. 2021;14(9):101162.

50. Levy BD, Clish CB, Schmidt B, Gronert K, Serhan CN. Lipid mediator class switching during acute inflammation: signals in resolution. Nat Immunol. 2001;2(7):612–619.

51. Chiang N, Dalli J, Colas RA, Serhan CN. Identification of resolvin D2 receptor mediating resolution of infections and organ protection. J Exp Med. 2015;212(8):1203–1217.

52. Kain V, Jadapalli JK, Tourki B, Halade GV. Inhibition of FPR2 impaired leukocytes recruitment and elicited non-resolving inflammation in acute heart failure. Pharmacol Res. 2019;146(104295.

53. Tourki B, Kain V, Pullen AB, et al. Lack of resolution sensor drives age-related cardiometabolic and cardiorenal defects and impedes inflammation-resolution in heart failure. Mol Metab. 2020;31(138-149.

54. Maddipati KR. Non-inflammatory Physiology of “Inflammatory” Mediators - Unalamation, a New Paradigm. Front Immunol. 2020;11(580117.

55. Hosseini Z, Marinello M, Decker C, et al. Resolvin D1 Enhances Necroptotic Cell Clearance Through Promoting Macrophage Fatty Acid Oxidation and Oxidative Phosphorylation. Arterioscler Thromb Vasc Biol. 2021;41(3):1062–1075.

56. Sulciner ML, Serhan CN, Gilligan MM, et al. Resolvins suppress tumor growth and enhance cancer therapy. J Exp Med. 2018;215(1):115–140.

57. Spite M, Norling LV, Summers L, et al. Resolvin D2 is a potent regulator of leukocytes and controls microbial sepsis. Nature. 2009;461(7268):1287-1291.

58. Groarke EM, Patel BA, Gutierrez-Rodrigues F, et al. Eltrombopag added to immunosuppression for children with treatment-naive severe aplastic anaemia. Br J Haematol. 2021;192(3):605–614.

59. Groarke EM, Feng X, Aggarwal N, et al. Efficacy of JAK1/2 inhibition in murine immune bone marrow failure. Blood. 2023;141(1):72–89.

60. Zeiser R, Polverelli N, Ram R, et al. Ruxolitinib for Glucocorticoid-Refractory Chronic Graft-versus-Host Disease. N Engl J Med. 2021;385(3):228–238.

61. Verstovsek S, Mesa RA, Gotlib J, et al. A double-blind, placebo-controlled trial of ruxolitinib for myelofibrosis. N Engl J Med. 2012;366(9):799–807.

