## Supplemental for "Resolvin E1 improves efferocytosis and rescues severe aplastic anemia in mice"

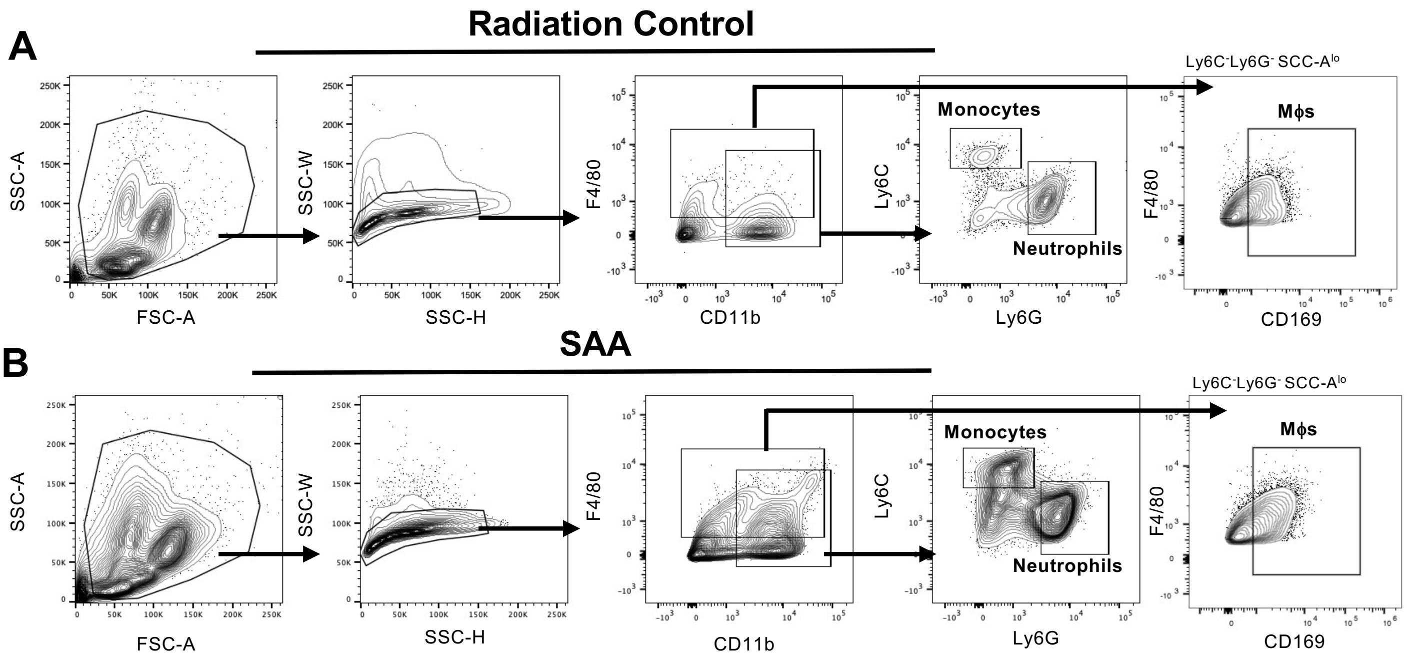


**Supplemental Figure 1. Myeloid gating strategy to identify bone marrow cells.** Representative flow cytometry analysis and gating for monocytes (CD11b$^{+}$ Ly6C$^{hi}$ Ly6G$^{-}$), neutrophils (CD11b$^{+}$ Ly6C$^{-}$ Ly6G$^{+}$), and macrophages (F480$^{+}$Ly6C$^{-}$Ly6G$^{-}$SSC-A$^{lo}$CD169$^{+}$) in bone marrow from radiation control (**A**) and SAA-induced (**B**) mice 10 dpst.

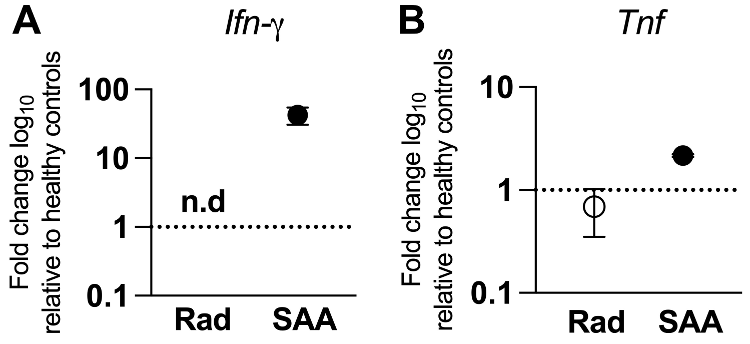


**Supplemental Figure 2. Gene expression in bone marrow monocytes**. Bone marrow monocytes ($CD11b^{+}Ly6C^{hi}Ly6G^{-}$) were sort-purified from radiation controls and SAA-induced mice on day 8 post-radiation and healthy (non-irradiated) control mice. RNA was extracted and cDNAs were generated for real time PCR analysis. Gene expression was normalized to β-actin and the $\log_{10}$ fold change, relative to healthy controls, is shown for *Ifn-γ* (**A**) and *Tnf* (**B**).

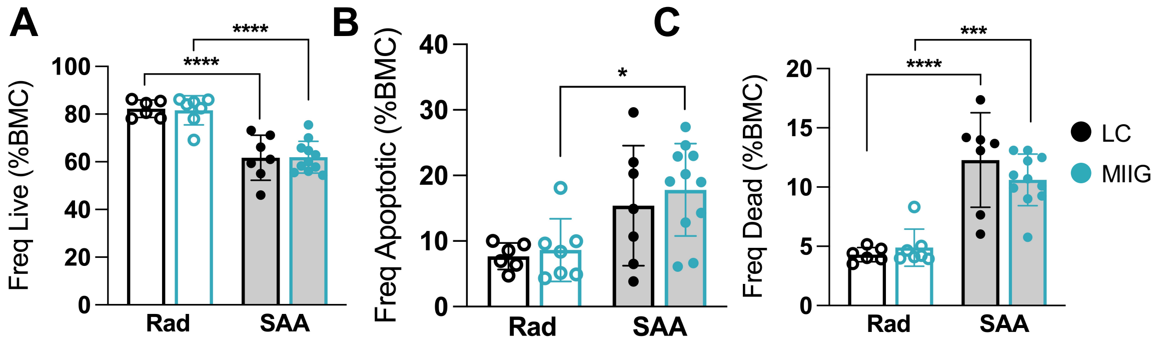


**Supplemental Figure 3. Apoptotic and dead bone marrow cells in the MIIG model.** F1 MIIG and littermate control (LC) mice were induced via the splenocyte transfer model and euthanized 10 days post-induction. Frequencies of live (AnnV$^-$ 7-AAD$^-$) (**A**), apoptotic (AnnV$^+$7-AAD$^-$) (**B**), and dead (7-AAD$^+$) (**C**) cells among total BM cells (%BMC). Data representative of two pooled independent experiments showing mean ± SD, n=6-11 per group. *p<0.05, *** p<0.001, **** p<0.0001

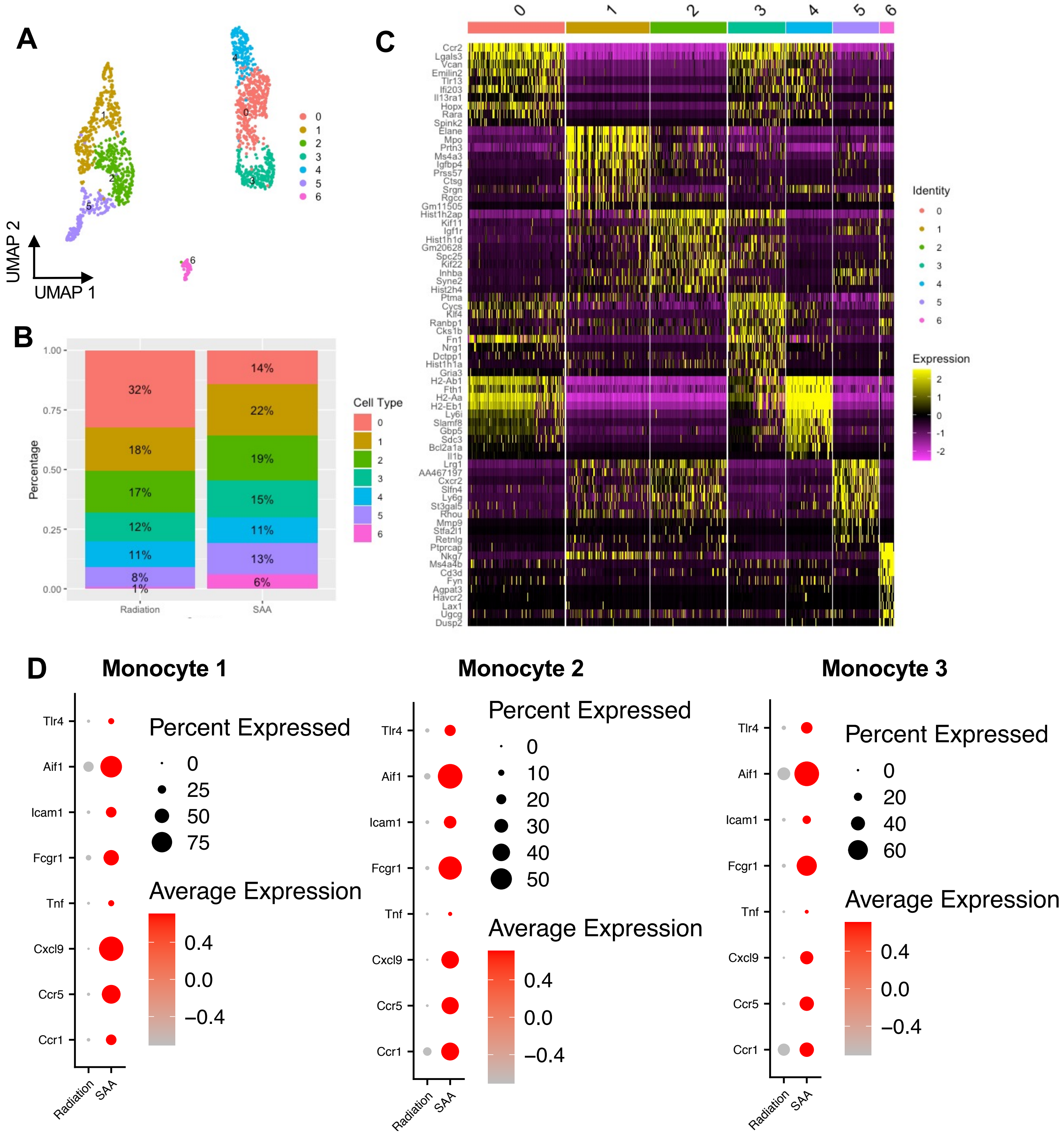


**Supplemental Figure 4. Single-cell analysis of bone marrow cells from radiation and SAA-induced mice.** (**A**) UMAP projection of flow sorted CD11b$^{+}$Ly6C$^{+}$Ly6G$^{-}$ bone marrow cells. A total of 6 clusters were identified and proportions (**B**) and marker genes (**C**) for each cluster is shown. (**D**) Dot plot of differentially expressed inflammatory genes in Monocyte 1 (cluster 4), Monocyte 2 (cluster 0), and Monocyte 3 (cluster 3) populations. The size of the dot corresponds to the percentage of cells expressing each gene, while the color represents the average gene expression level.

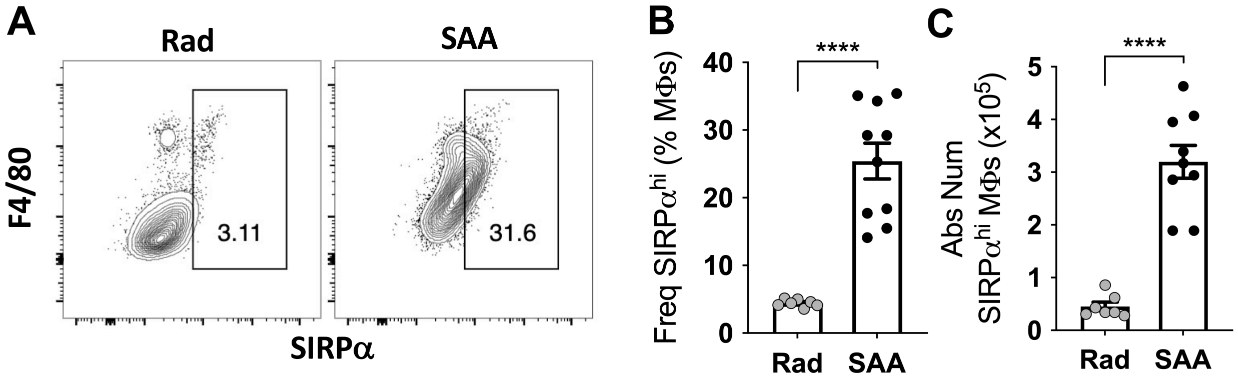


**Supplementary Figure 5. SIRPα expression increases on macrophages during SAA.** F1 hybrid mice were induced to develop SAA and BM harvested 10 dpst. (**A**) Representative gating for SIRPα$^{hi}$ macrophages. The frequency (**B**) and absolute number (**C**) of SIRPα$^{hi}$ macrophages. Data representative of two experiments showing mean ± SD, n=7-10 per group, significance was determined using a Student's t-test. **** $p<0.0001$

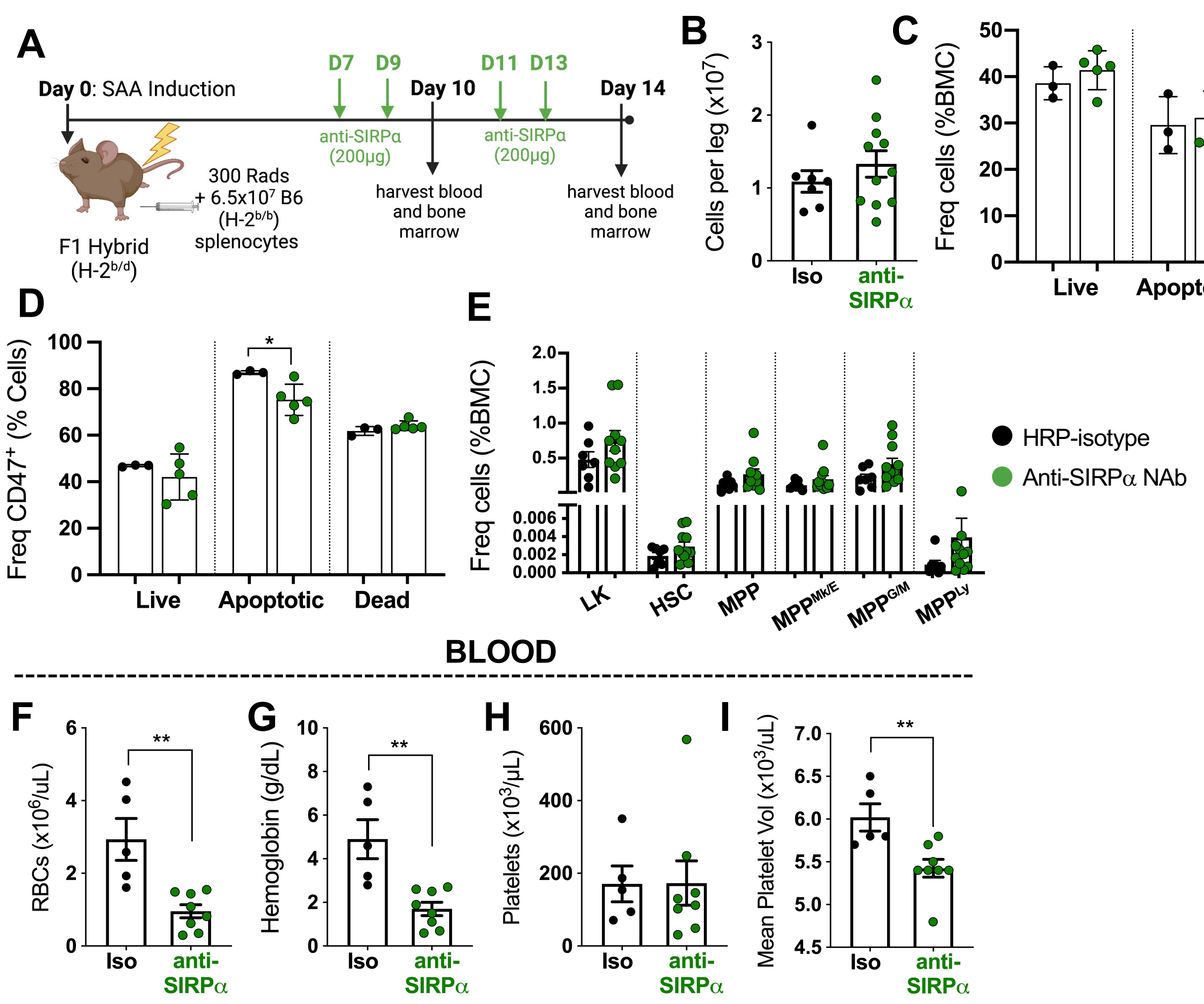


**Figure 6. Anti-SIRPa treatment reduces apoptotic and dead cells but exacerbates anemia during SAA.** (**A**) F1 mice were treated with anti-SIRPα neutralizing antibody (clone P84; 200mg) days 7, 9, 11, and 13 post SAA-induction. BM and blood was harvested 10 (**B-E**) or 14 dpst (**F-I**). (**B**) Absolute number of bone marrow cells per leg. (**C**) Frequencies of live, apoptotic, and dead cells as a percent of total BM cells (%BMC). (**D**) CD47 expression on live, apoptotic, and dead BM cells. (**E**) Frequencies of HSPC populations as a percent of total BM cells. Total RBCs (**F**), hemoglobin (**G**), platelets (**H**), and mean platelet volume (**I**). Data from one independent experiment per timepoint showing mean ± SD, n=3-8 per group. Significance was determined using a Student's t-test for **B** and **F-I**; Two-way ANOVA with Tukey's multiple comparison was done for **C-E**. *p<0.05, ** p<0.01

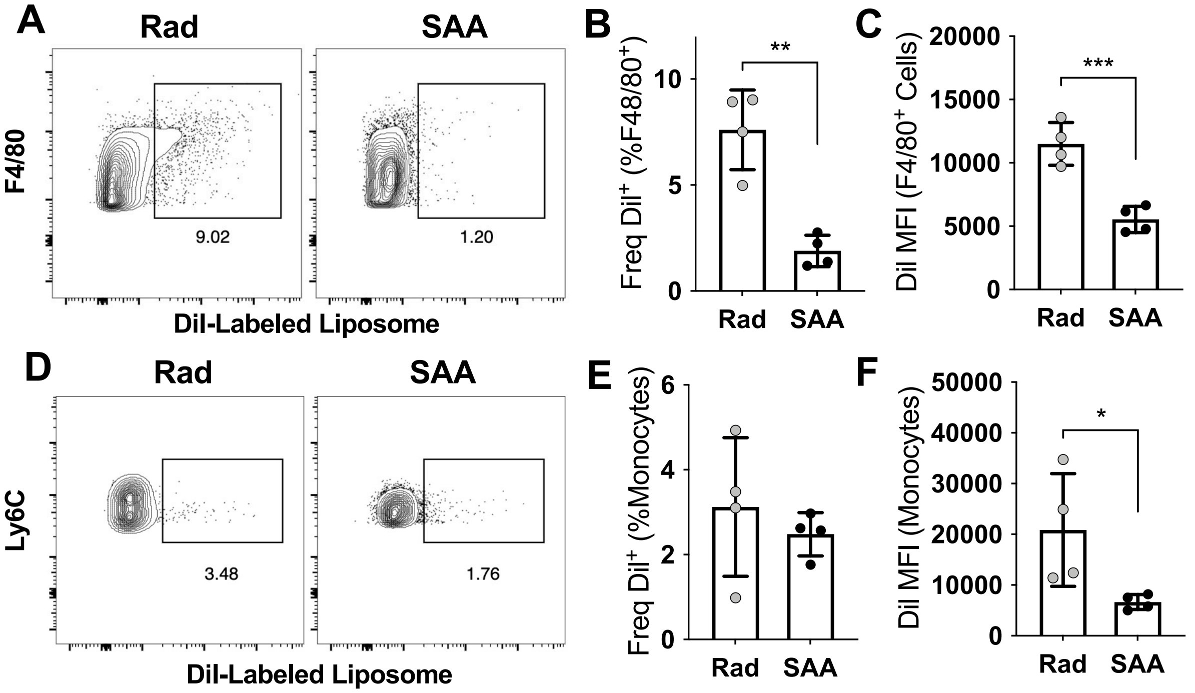


**Supplementary Figure 7. Phagocytosis is impaired during SAA.** F1 hybrid mice were induced via the splenocyte transfer model and administered 200μL of fluorescent (DiI-labeled) or control liposomes via retro-orbital (I.V.) injection 9 dpst; BM was harvested 10 dpst. Representative staining, frequency, and MFI of DiI-labeled liposomes on F4/80⁺ macrophages (**A-C**) or monocytes (**D-F**). Data from one experiment showing mean ± SD, n=4 per group. Significance was determined using a Student's t-test. * p<0.05, ** p<0.01, *** p<0.001

**A**

| Variable | PC1 | PC2 |
|---|---|---|
| 5(S), 15(S)-DiHETE | -0.7550 | 0.5998 |
| 5(S), 12(S)-DiHETE | 0.0580 | 0.5027 |
| 5(S), 6(R)-DiHETE | -0.4619 | 0.8143 |
| LTB4 | 0.1632 | -0.3811 |
| LXA4 | 0.8858 | 0.2824 |
| Maresin 1 | 0.8594 | 0.4294 |
| 5-HEPE | 0.9178 | -0.0869 |
| 11-HEPE | -0.4755 | 0.7536 |
| 12-HEPE | 0.8695 | 0.4305 |
| 15(S)-HEPE | 0.7880 | 0.5750 |
| 18-HEPE | 0.8882 | 0.4015 |
| 5-HETE | 0.5508 | 0.1831 |
| 11-HETE | 0.3563 | 0.7878 |
| 12-HETE | 0.9079 | 0.3905 |
| 15-HETE | 0.9520 | 0.2854 |
| 4-HDoHE | 0.4563 | -0.6747 |
| 7-HDoHE | 0.9488 | 0.1933 |
| 14-HDoHE | -0.8120 | 0.3061 |
| 13-HDoHE | -0.6562 | 0.5747 |
| 17-HDoHE | -0.8143 | 0.3175 |
| 10S,17S-DiHDoHE | 0.4397 | 0.4007 |
| PD1 | -0.2407 | 0.3740 |
| 12(S)-HHTrE | -0.8749 | 0.2765 |
| PGE2 | 0.3089 | 0.9263 |
| PGD2 | -0.4532 | 0.8276 |
| PGF2a | -0.1264 | -0.8422 |
| TXB2 | -0.4281 | 0.8502 |
| RvD6 | -0.7550 | 0.5998 |
| RvD5 | 0.0580 | 0.5027 |

**B**

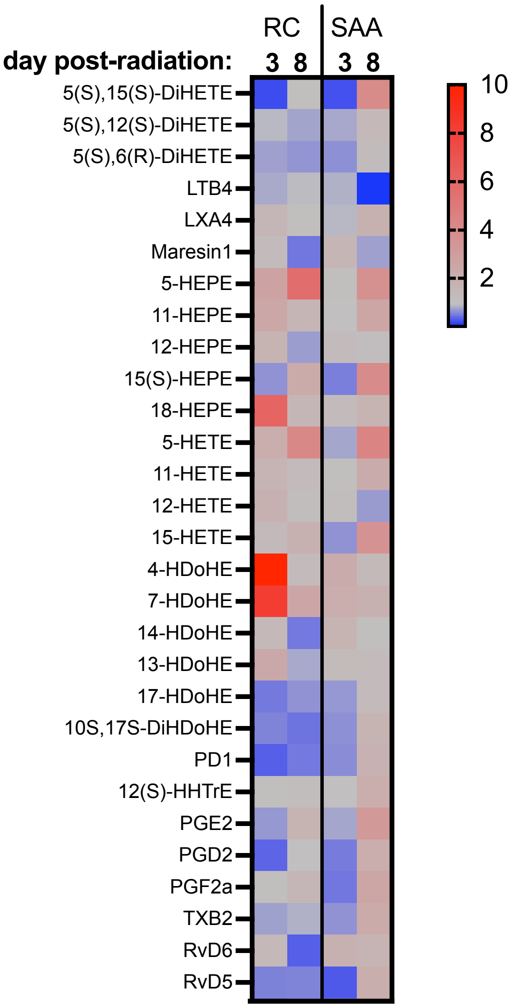


**Supplemental Figure 8. Lipid mediator analysis of the bone marrow.** Table of the loadings for PC1 and PC2 for data in PC scores graph in Figure 5A (**A**). Heat map of lipid mediators detected in bone marrow from radiation controls or SAA-induced mice on day 3 or 8 post-radiation. Data show fold change relative to healthy mice (n=2-4 mice per group) (**B**) .

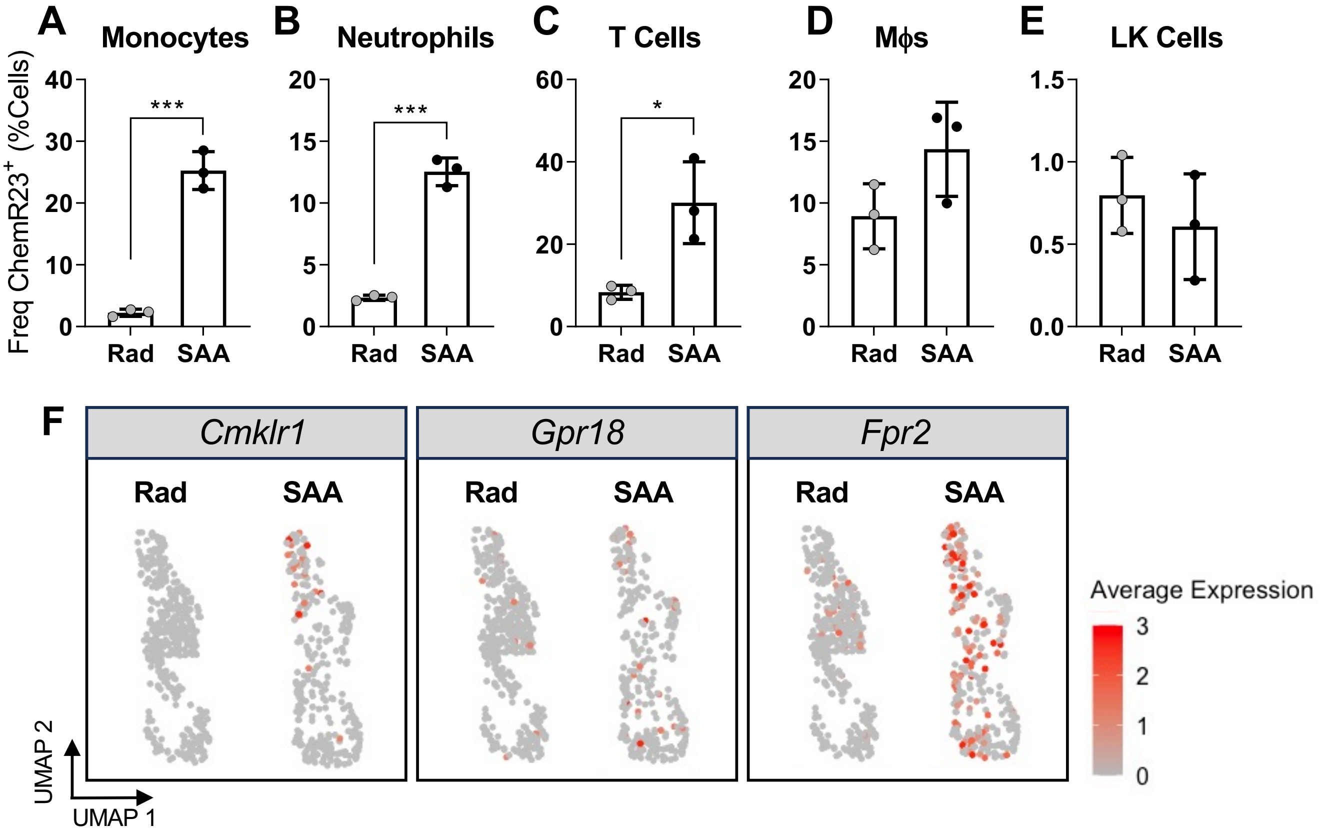


**Supplemental Figure 9. ChemR23 expression increases on myeloid cells and T cells during SAA.** The expression of ChemR23 increases on monocytes (**A**), neutrophils (**B**), and T cells **(C)** in SAA mice, compared to radiation control 10 dpst. The frequency of ChemR23 does not significantly change on macrophages (**D**) or total LK cells (**E**) during SAA. Data from one independent experiment showing mean ± SD, n=3 per group. Significance was determined using a Student's t-test. * $p<0.05$, *** $p<0.001$. (**F**) Gene expression data of SPM receptors from single-cell RNA sequencing of BM monocytes.

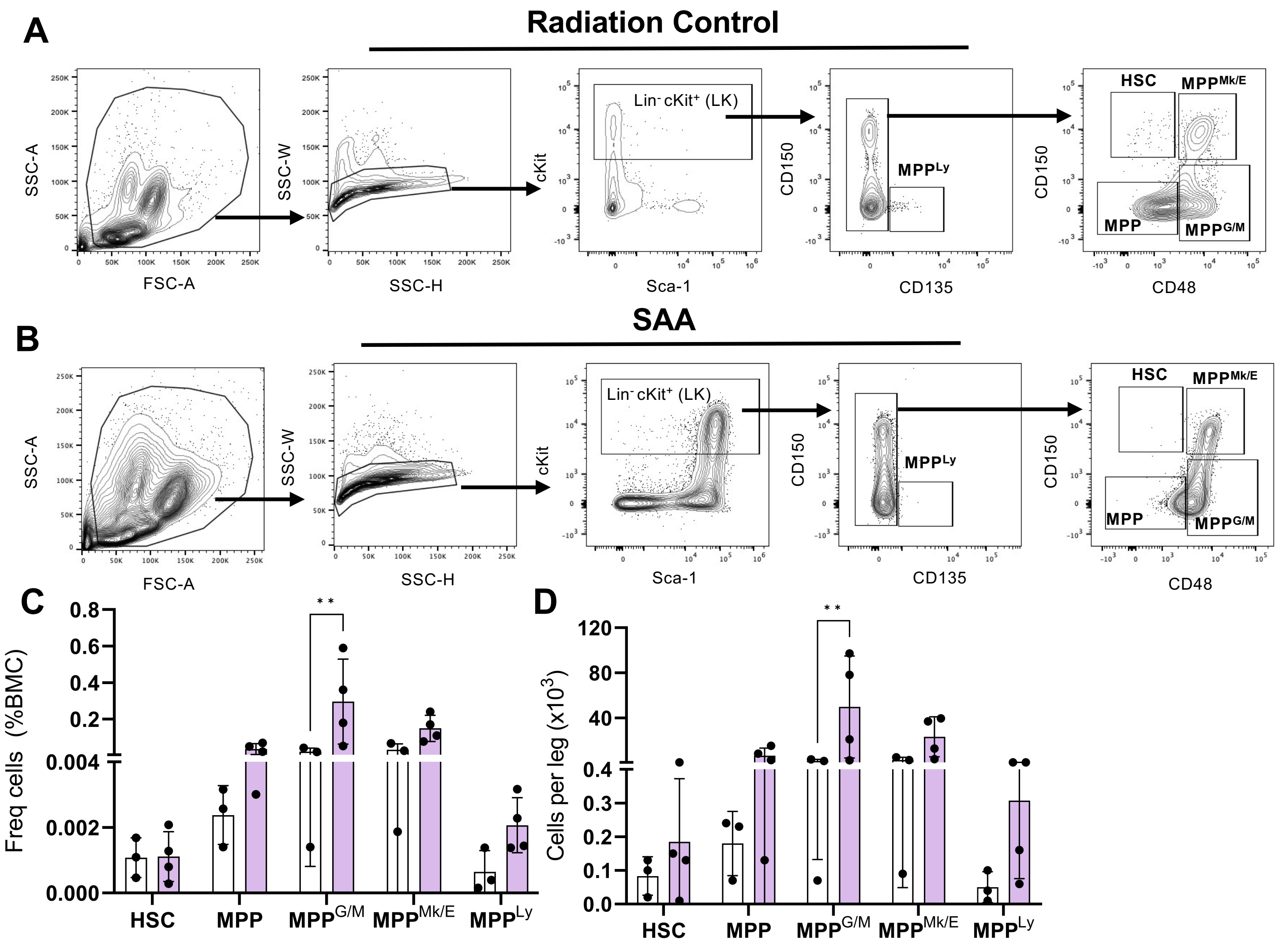


**Supplemental Figure 10. HSPC populations expand with RvE1 treatment.** Gating strategy for identifying HSCs (CD48⁻CD150⁺), MPPs (CD48⁻CD150⁻), MPP$^{Mk/E}$(CD48⁺CD150⁺), and MPP$^{G/M}$ (CD48⁺CD150⁻) subpopulations among Lineage-negative (Lin⁻) cKit⁺ bone marrow cells from radiation control (**A**) and SAA (**B**) mice 10 dpst. SAA mice were induced and treated with 250ng RvE1 days 7, 9, and 11 post-induction; BM harvested 12 dpst. Frequency (**C**) and absolute number (**D**) of HSCs, MPPs, MPP$^{Mk/E}$, and MPP$^{G/M}$ in the BM. Significance was determined using Two-way ANOVA with Tukey's multiple comparison test. ** $p<0.01$

| Antigen | Clone | Vendor | Catalog Number (Fluor) |
|---|---|---|---|
| 7AAD | - | BioLegend | 420403 (PerCP Cy5.5) |
| Annexin V | - | BioLegend | 640924 (BV421) |
| CD11b | M1/70 | BioLegend | 101206 (FITC), 101216 (PE Cy7) |
| CD135 | AF210 | eBioscience | 135308 (APC) |
| CD150 | TC15-12F12.2 | BioLegend | 115941 (BV711) |
| CD169 | 3D6.112 | BioLegend | 142411 (PE Cy7) |
| CD38 | HM48-1 | BioLegend | 103418 (Pacific Blue) |
| CD47 | Miap301 | BioLegend | 127507 (PE), 127515 (PerCP Cy5.5) |
| ChemR23 | 477806 | R&D Systems | FAB7610P (PE) |
| cKit | 2B8 | BioLegend | 105812 (APC) |
| F4/80 | CI:A3-1 | Abcam | AB105080 (APC) |
| Ly6C | HK1.4 | BioLegend | 128033 (BV510), 128013 (Pacific Blue) |
| Ly6G | 1A8 | BioLegend | 127639 (BV605) |
| MerTK | Polyclonal | R&D Systems | BAF591 (Biotinylated) |
| Sca-1 | D7 | BioLegend | 108114 (PE Cy7) |
| SIRP$\alpha$ | P84 | BioLegend | 144012 (PE), 144013 (APC) |

**Supplementary Table 1. Antibody Information.** Each antigen and respective clone used for flow cytometry analysis.

| Lipid | Healthy | Day 3 (RC) | Day 3 (SAA) | Day 8 (RC) | Day 8 (SAA) |
|---|---|---|---|---|---|
| 5(S),15(S)-DiHETE | 0.33 ± 0.12 | 0.09 ± 0.02 | 0.10 ± 0.03 | 0.37 ± 0.16 | 1.34 ± 0.57 |
| 5(S),12(S)-DiHETE | 8.49 ± 1.80 | 8.11 ± 2.00 | 7.08 ± 1.11 | 6.92 ± 0.52 | 12.08 ± 0.55 |
| 5(S),6(R)-DiHETE | 2.75 ± 0.18 | 2.15 ± 0.43 | 1.88 ± 0.19 | 1.96 ± 0.17 | 3.43 ± 0.30 |
| Maresin1 | 0.15 ± 0.02 | 0.19 ± 0.08 | 0.25 ± 0.05 | 0.08 | 0.12 ± 0.05 |
| 5-HEPE | 0.09 ± 0.06 | 0.25 ± 0.02 | 0.10 ± 0.03 | 0.52 ± 0.21 | 0.35 ± 0.21 |
| 11-HEPE | 0.53 ± 0.18 | 1.26 ± 0.25 | 0.54 ± 0.12 | 0.86 ± 0.43 | 1.34 ± 0.40 |
| 12-HEPE | 64.57 ± 10.47 | 109.05 ± 15.23 | 83.16 ± 4.25 | 49.31 ± 8.19 | 74.61 ± 6.53 |
| 15(S)-HEPE | 2.99 ± 0.91 | 2.08 ± 0.67 | 1.72 ± 0.61 | 6.66 ± 4.11 | 12.22 ± 5.28 |
| 18-HEPE | 0.06 ± 0.02 | 0.40 ± 0.12 | 0.08 ± 0.02 | 0.10 ± 0.06 | 0.11 ± 0.03 |
| 5-HETE | 1.37 ± 0.31 | 2.92 ± 0.76 | 1.13 ± 0.37 | 5.84 ± 1.92 | 6.06 ± 2.07 |
| 11-HETE | 17.87 ± 2.48 | 30.01 ± 2.73 | 19.19 ± 2.23 | 22.46 ± 4.82 | 39.38 ± 8.13 |
| 12-HETE | 485.44 ± 311.84 | 938.49 ± 487.50 | 583.03 ± 216.83 | 572.03 ± 292.68 | 366.19 ± 32.49 |
| 15-HETE | 8.16 ± 3.39 | 11.14 ± 3.11 | 5.74 ± 0.66 | 15.49 ± 4.01 | 29.70 ± 10.44 |
| 4-HDoHE | 0.14 ± 0.05 | 2.17 ± 0.40 | 0.30 ± 0.08 | 0.18 ± 0.07 | 0.19 ± 0.06 |
| 7-HDoHE | 0.08 ± 0.05 | 0.68 ± 0.07 | 0.18 ± 0.07 | 0.21 ± 0.04 | 0.15 ± 0.09 |
| 14-HDoHE | 42.80 ± 6.45 | 61.35 ± 8.66 | 72.77 ± 7.10 | 22.81 ± 7.88 | 49.05 ± 9.43 |
| 13-HDoHE | 2.78 ± 0.68 | 6.51 ± 0.85 | 3.75 ± 0.64 | 2.34 ± 0.75 | 3.53 ± 0.98 |
| 17-HDoHE | 12.91 ± 0.49 | 7.06 ± 1.81 | 9.39 ± 2.16 | 9.00 ± 3.73 | 16.79 ± 5.40 |
| 10S,17S-DiHDoHE | 2.44 ± 0.58 | 1.49 ± 0.64 | 1.66 ± 0.50 | 1.23 ± 0.87 | 4.25 ± 2.25 |
| PD1 | 0.54 ± 0.15 | 0.20 ± 0.08 | 0.36 ± 0.17 | 0.29 ± 0.20 | 0.99 ± 0.59 |
| 12(S)-HHTrE | 6.94 ± 1.04 | 7.74 ± 0.43 | 7.01 ± 0.69 | 8.20 ± 0.41 | 14.54 ± 1.30 |
| PGE2 | 12.54 ± 4.97 | 9.24 ± 3.60 | 10.36 ± 2.49 | 21.71 ± 5.69 | 41.15 ± 10.62 |
| PGD2 | 32.16 ± 3.54 | 13.20 ± 1.06 | 18.13 ± 1.33 | 33.32 ± 11.67 | 68.53 ± 15.81 |
| PGF2a | 1.35 ± 0.09 | 1.53 ± 0.28 | 0.71 ± 0.14 | 2.08 ± 0.69 | 3.51 ± 0.69 |
| TXB2 | 20.27 ± 2.03 | 16.17 ± 1.70 | 14.19 ± 1.83 | 18.21 ± 1.65 | 45.21 ± 7.91 |
| RvD6 | 0.16 ± 0.07 | 0.24 ± 0.03 | 0.32 ± 0.09 | 0.06 ± 0.00 | 0.27 ± 0.16 |
| RvD5 | 0.56 ± 0.19 | 0.33 ± 0.12 | 0.19 ± 0.09 | 0.34 ± 0.22 | 1.13 ± 0.58 |

**Supplementary Table 2. Quantification of lipids in bone marrow.** Concentrations of the indicated lipid in bone marrow from healthy, radiation control (RC), and SAA-induced mice at the indicated time post-radiation. Concentrations are pg per mg total protein.
